# The genome of *Trichoplusia ni*, an agricultural pest and novel model for small RNA biology

**DOI:** 10.1101/183921

**Authors:** Yu Fu, Yujing Yang, Han Zhang, Gwen Farley, Junling Wang, Kaycee A. Quarles, Zhiping Weng, Phillip D. Zamore

## Abstract

The cabbage looper, *Trichoplusia ni* (Lepidoptera: Noctuidae), is a destructive insect pest that feeds on a wide range of plants. The High Five cell line (Hi5), originally derived from *T. ni* ovaries, is often used for efficient expression of recombinant proteins. Here, we report a draft assembly of the 368.2 Mb *T. ni* genome, with 90.6% of all bases assigned to one of its 28 chromosomes and predicted 14,037 predicted protein-coding genes. Manual curation of gene families involved in chemoreception and detoxification reveals *T. ni*-specific gene expansions that may explain its widespread distribution and rapid adaptation to insecticides. Using male and female genome sequences, we define Z-linked and repeat-rich W-linked sequences. Transcriptome and small RNA data from *T. ni* thorax, ovary, testis, and Hi5 cells reveal distinct expression profiles for 295 microRNA- and >393 piRNA-producing loci, as well as 39 genes encoding core small RNA pathway proteins. siRNAs target both endogenous transposons and the exogenous TNCL virus. Surprisingly, *T. ni* siRNAs are not 2´-*O*-methylated. Five piRNA-producing loci account for 34.9% piRNAs in the ovary, 49.3% piRNAs in the testis, and 44.0% piRNAs in Hi5 cells. Nearly all of the W chromosome is devoted to piRNA production: >76.0% of bases in the assembled W produce piRNAs in ovary. To enable use of the *T. ni* germline-derived Hi5 cell line as a model system, we have established efficient genome editing and single-cell cloning protocols. Taken together, the *T. ni* genome provides insights into pest control and allows Hi5 cells to become a new tool for studying small RNAs ex vivo.

## Introduction

Lepidoptera (moths and butterflies), one of the most species-rich orders of insects, comprises more than 170,000 known species (*Mallet J, 2007*;), many of which are agricultural pests. One of the largest lepidopteran families, Noctuidae, diverged over 100 million years ago (mya) from the Bombycidae—best-known for the silkworm, *Bombyx mori* (*Rainford JL et al., 2014*). The Noctuidae family member cabbage looper (*Trichoplusia ni*) is a widely distributed generalist pest that feeds on cruciferous crops such as broccoli, cabbage, and cauliflower (). *T. ni* has evolved resistance to the chemical insecticide Dichlorodiphenyltrichloroethane (DDT; [*McEwen FL and Hervey GER, 1956*]) and the biological insecticide *Bacillus thuringiensis* toxin (*Janmaat AFMyers J, 2003*), rendering pest control increasingly difficult. A molecular understanding of insecticide resistance requires a high-quality *T. ni* genome and transcriptome.

Hi5 cells derive from *T. ni* ovarian germ cells (*Granados RR et al., 1986*; *Granados RR et al., 1994*). Hi5 cells are a mainstay of recombinant protein production using baculoviral vectors (*Wickham TJ et al., 1992*) and hold promise for the commercial-scale production of recombinant adeno-associated virus for human gene therapy (*Kotin RM, 2011*; *van Oers MM et al., 2015*). Hi5 cells produce abundant microRNAs (miRNAs), small interfering RNAs (siRNAs), and PIWI-interacting RNAs (*Kawaoka S et al., 2009*) (piRNAs), making them one of just a few cell lines suitable for the study of all three types of animal small RNAs. The most diverse class of small RNAs, piRNAs protect the genome of animal reproductive cells by silencing transposons (*Brennecke J et al., 2007*; *Reuter M et al., 2011*). The piRNA pathway has been extensively studied in the dipteran insect *Drosophila melanogaster* (fruit fly), but no piRNA-producing, cultured cell lines exist for dipteran germline cells. *T. ni* Hi5 cells grow rapidly without added hemolymph (*Hink WF, 1970*), are readily transfected, and— unlike the more difficult to grow *B. mori* BmN4 cells (*Iwanaga M et al., 2014*), which also express germline piRNAs—remain homogeneously undifferentiated even after prolonged culture. In contrast to *B. mori*, no *T. ni* genome sequence is available, limiting the utility of Hi5 cells.

To further understand this agricultural pest and its Hi5 cell line, we combined divers genomic sequencing data to assemble a chromosome-level, high-quality *T. ni* genome. Half the genome sequence resides in scaffolds >14.2 megabases (Mb), and >90% is assembled into 28 chromosome-length scaffolds. Automated gene prediction and subsequent manual curation, aided by extensive RNA-seq data, allowed us to examine gene orthology, gene families such as detoxification proteins, sex determination genes, and the miRNA, siRNA, and piRNA pathways. Our data allowed assembly of the gene-poor, repeat-rich W chromosome, which remarkably produces piRNAs across most of its length. To enable the use of cultured *T. ni* Hi5 cells as a novel insect model system, we established methods for efficient genome editing using the CRISPR/Cas9 system (*Ran FA et al., 2013*) as well as single-cell cloning. With these new tools, *T. ni* promises to become a powerful companion to flies to study gene expression, small RNA biogenesis and function, and mechanisms of insecticide resistance in vivo and in cultured cells.

## Results

### Genome sequencing and assembly

We combined Pacific Biosciences long reads and Illumina short reads (Figure 1A, Table 1, and Materials and methods) to sequence genomic DNA from Hi5 cells and *T. ni* male and female pupae. The initial genome assembly from long reads (46.4× coverage with reads >5 kb) was polished using paired-end (172.7× coverage) and mate-pair reads (172.0× coverage) to generate 1,976 contigs spanning 368.2 megabases (Mb). Half of genomic bases reside in contigs >621.9 kb (N50). Hi-C long-range scaffolding (186.5× coverage) produced 1,031 scaffolds (N50 = 14.2 Mb), with >90% of the sequences assembled into 28 major scaffolds. Karyotyping of metaphase Hi5 cells revealed that these cells have 112 ± 5 chromosomes (Figure 1B, Figure 1–figure supplement 1). Because lepidopteran cell lines are typically tetraploid (), we conclude that the ~368.2 Mb *T. ni* genome comprises 28 chromosomes: 26 autosomes plus W and Z sex chromosomes (see below).

**Table 1.** Genome and gene set statistics for *T. ni* and *B. mori (The ISG, 2008)*. Cytochrome P45Os, glutathione S-transferases, carboxylesterases, and ATP-binding cassette transporters for *B. mori* were retrieved from *(Yu Q et al., 2008; Yu Q-Y et al., 2009; Ai J et al., 2011; Liu S et al., 2011)*.

|  | *T. ni* | *B. mori* |
| --- | --- | --- |
| **Genome Metrics** |  |  |
| Genome size (Mb) | 368.2 | 431.7 |
| Chromosome count | 28 | 28 |
| Scaffold N50 (Mb) | 14.2 | 3.7 |
| Contig N50 (Kb) | 621.9 | 15.5 |
| Mitochondrial genome (Kb) | 15.8 | 15.7 |
| **Quality Control Metrics** |  |  |
| BUSCO complete (%) | 97.5 | 95.5 |
| CRP genes (%) | 100% | 100% |
| OXPHOS genes (%) | 100% | 100% |
| **Genomic Features** |  |  |
| Repeat content (%) | 20.5% | 43.6% |
| GC content | 35.6% | 37.3% |
| CpG (O/E) | 1.07 | 1.13 |
| Coding (%) | 5.58 | 4.11 |
| Sex chromosomes | ZW | ZW |
| **Gene Statistics** |  |  |
| Protein-coding genes | 14,043 | 14,623 |
| with Pfam matches | 9,295 | 9,685 |
| with GO terms | 9,790 | 10,148 |
| Cytochrome P450 proteins | 108 | 83 |
| Glutathione S-transferases | 34 | 23 |
| Carboxylesterases | 87 | 76 |
| ATP-binding cassette transporters | 54 | 51 |
| Universal orthologs lost | 156 | 75 |
| Species-specific genes | 3,098 | 2,313 |

**Figure 1.**
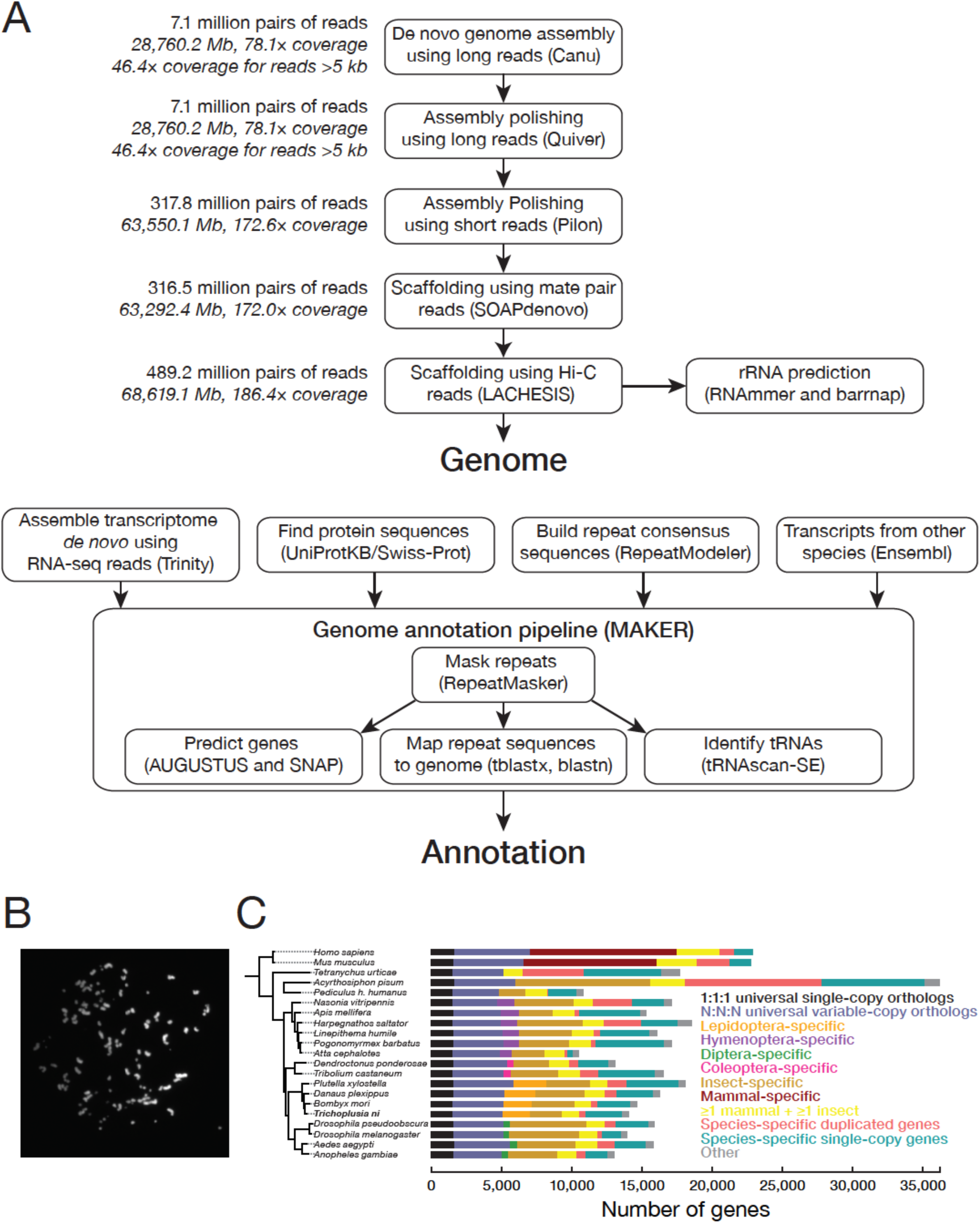
Chromosomes and genes in the *T. ni* genome. (A) Genome assembly and annotation workflow. (B) An example of a DAPI-stained spread of Hi5 cell mitotic chromosomes used to determine the karyotype. (C) Phylogenetic tree and orthology assignment of *T. ni* with 18 arthropod and two mammalian genomes. Colors denote gene categories. The category 1:1:1 represents universal single-copy orthologs, allowing absence and/or duplication in one genome. N:N:N orthologs include orthologs with variable copy numbers across species, allowing absence in one genome or two genomes from different orders. Lepidoptera-specific genes are present in at least three of the four lepidopteran genomes; Hymenoptera-specific genes are present in at least one wasp or bee genome and at least one ant genome. Coleoptera-specific genes are present in both coleopteran genomes; Diptera-specific genes present in at least one fly genome and one mosquito genome. Insect indicates other insect-specific genes. Mammal-specific genes are present in both mammalian genomes. The phylogenetic tree is based on the alignment of 1:1:1 orthologs.

To evaluate the completeness of the assembled *T. ni* genome, we compared it to the Arthropoda data set of the Benchmark of Universal Single-Copy Orthologs (*Simão FA et al., 2015*) (BUSCO v3). The *T. ni* genome assembly captures 97.5% of these gene orthologs, more than either the silkworm (95.5%) or monarch butterfly (*D. plexippus*; 97.0%) genomes (Supplementary file 1A). All 79 ribosomal proteins conserved between mammals and *D. melanogaster* (*Yoshihama M et al., 2002*; *Marygold SJ et al., 2007*) have orthologs in *T. ni*, further evidence of the completeness of the genome assembly (Supplementary file 1B). Finally, a search for genes in the highly conserved nuclear oxidative phosphorylation (OXPHOS) pathway (*Porcelli D et al., 2007*) uncovered *T. ni* orthologs for all known *D. melanogaster* OXPHOS genes (Supplementary file 1C).

We also de novo assembled the *T. ni* genomes using paired-end DNA-seq data obtained from male and female pupae respectively, but the resulting assemblies are fragmented (scaffold N50 ≤2.4 kb, Supplementary file 1D), likely due to limitations of short-insert libraries and high levels of heterozygosity commonly observed for insect genomes (*Keeling CI et al., 2013*; *You M et al., 2013*). Comparison of the animal genome contigs with the Hi5 genome as the reference shows high concordance (≤ 1.37% of animal contigs are misassembled, Supplementary file 1D), indicating that our Hi5 genome assembly is representative of the *T. ni* animal genome.

### Gene orthology

We annotated 14,034 protein-coding genes in the *T. ni* genome, similar to other Lepidoptera (*Challis RJ et al., 2016*) (Supplementary file 1E). We analyzed the homology of *T. ni* genes to genes in 20 species that span the four common insect orders (Lepidoptera, Diptera, Coleoptera, Hymenoptera), non-insect arthropods, and mammals (Figure 1C, Figure 1–figure supplement 2). These 21 species define 30,448 orthology groups each containing orthologous proteins from two or more species (*Hirose Y and Manley JL, 1997*); 9,112 groups contain at least one *T. ni* gene. In all, 10,936 *T. ni* protein-coding genes are orthologous to at least one gene among the 20 reference species.

*T. ni* contains 2,287 Lepidoptera-specific (*T. ni, B. mori, D. plexippus*, and *P. xylostella* [diamondback moth]) orthology groups. Far fewer orthology groups are unique to Diptera (404), Coleoptera (371), or Hymenoptera (1,344), suggesting that the lepidopteran lifestyle requires more order-specific genes. The *T. ni* genome additionally contains 3,098 orphan protein-coding genes for which we could detect no orthologous sequences in the 20 reference species. Of these orphan genes, 14.5% are present as two or more copies in the genome (“in-paralogs”), suggesting they evolved recently; some of these in-paralogs may have arisen by gene duplication after the divergence of *T. ni* and *B. mori* ~111 mya (*Gaunt MW and Miles MA, 2002*; *Rota-Stabelli O et al., 2013*; *Wheat CW* and *Wahlberg N, 2013*; *Rainford JL et al., 2014*).

### Opsins

The ability of insects to respond to light is crucial to their survival. Opsins, members of the G-protein-coupled receptor superfamily, play important roles in vision. Covalently bound to light-sensing chromophores, opsins absorb photons and activate the downstream visual transduction cascade (*Terakita A, 2005*). The *T. ni* genome encodes ultraviolet, blue, and long-wavelength opsins, suggesting that, despite being a nocturnal insect, *T. ni* retains the full repertoire of insect opsins and has color vision (*Zimyanin VL et al., 2008*) (Figure 1–figure supplement 3). *T. ni* also encodes an ortholog of the non-visual Rh7 opsin, which is found in a variety of insects (*Initiative IGG, 2014*; *Futahashi R et al., 2015*) and participates in the entrainment of circadian rhythms by sensing violet light in the brain of *D. melanogaster* (*Ni JD et al., 2017*), and an ortholog of the vertebrate-like opsin, pterosin, which was first detected in honeybee (*A. mellifera*) brain and is found widely among insects except for *Drosophilid* flies (*Velarde RA et al., 2005*).

### Sex determination

Understanding the *T. ni* sex-determination pathway holds promise for engineering sterile animals for pest management. ZW and ZO chromosome systems determine sex in lepidopterans: males are ZZ and females are either ZW or ZO (*Traut W et al., 2007*). To determine which system *T. ni* uses and identify which contigs belong to the sex chromosomes, we sequenced genomic DNA from male and female pupae and calculated the male:female coverage ratio for each contig (Materials and methods). We found that 175 presumably Z-linked contigs (20.0 Mb) had approximately twice the coverage in male compared to female DNA (median male:female ratio = 1.92; Figure 2A, Figure 2–figure supplement 1A). Another 276 contigs (11.1 Mb) had low coverage in males (median male:female ratio = 0.111), suggesting they are W-linked. We conclude that sex is determined in *T. ni* by a ZW system in which males are homogametic (ZZ) and females are heterogametic (ZW).

**Figure 2.**
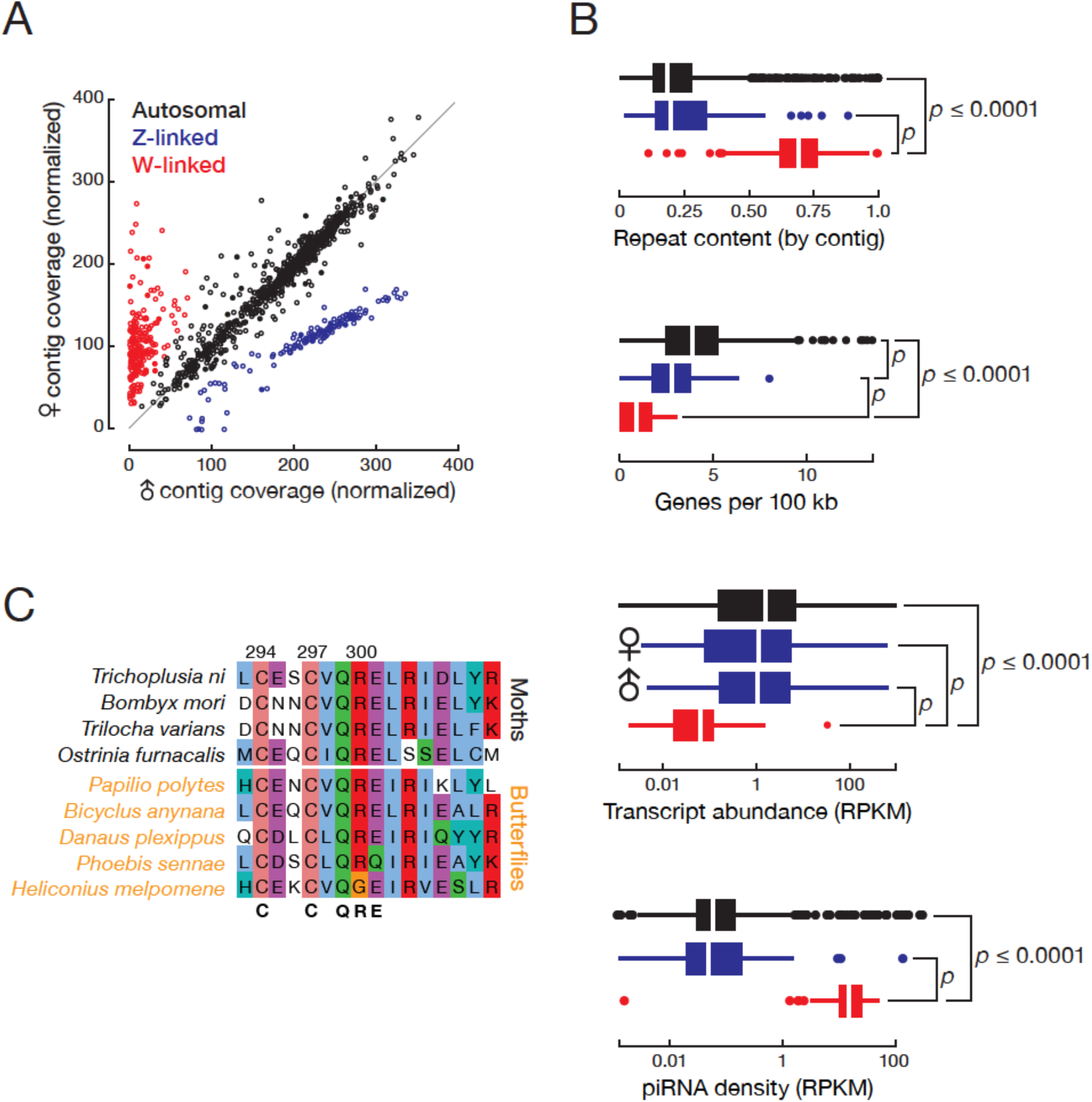
*T. ni* males are ZZ and females are ZW. (A) Normalized contig coverage in males and females. (B) Relative repeat content, gene density, transcript abundance (female and male thoraces), and piRNA density of autosomal, Z-linked, and W-linked contigs (ovary). (C) Multiple sequence alignment of the conserved region of the sex determining gene *masc* among the lepidopteran species.

For some lepidopteran species, dosage compensation has been reported to equalize Z-linked transcript abundance between ZW females and ZZ males in the soma, while other species show higher expression of Z-linked genes in males (*Walters JR et al., 2015*; *Gu L et al., 2017*). In the soma, *T. ni* compensates for Z chromosome dosage: transcripts from Z-linked genes are approximately equal in male and female thoraces (Z ≈ ZZ, Figure 2B). In theory, somatic dosage compensation could reflect increased transcription of the single female Z chromosome, reduced transcription of both male Z chromosomes, or silencing of one of the two male Z chromosomes.

To distinguish among these possibilities, we compared the abundance of Z-linked and autosomal transcripts (Z/AA in female and ZZ/AA in male, Figure 2–figure supplement 1B and 1C). Z-linked transcripts in the male thorax are expressed at lower levels than autosomal transcripts, but not as low as half (ZZ ≈ 70% AA). These data support a dosage compensation mechanism that decreases transcription from each Z chromosome in the *T. ni* male soma, but does not fully equalize Z-linked transcript levels between the sexes (Z ≈ ZZ ≈ 70% AA). In contrast, *T. ni* lacks germline dosage compensation: in the ovary, Z-linked transcript abundance is half that of autosomal transcripts (Z ≈ 50% AA), whereas in testis, Z-linked and autosomal transcripts have equal abundance (ZZ ≈ AA). We conclude that *T. ni*, like *B. mori* (*Walters JR and Hardcastle TJ, 2011*), *Cydia pomonella* (*Gu L et al., 2017*), and *Heliconius* butterflies (*Walters JR et al., 2015*), compensates for Z chromosome dosage in the soma by reducing gene expression in males, but does not decrease Z-linked gene expression in germline tissues.

Little is known about lepidopteran W chromosomes. The W chromosome is not included in the genome assemblies of *Manduca sexta* (*Kanost MR et al., 2016*) or *B. mori* (*The ISG, 2008*), and earlier efforts to assemble the silkworm W resulted in fragmented sequences containing transposons (*Abe H et al., 2005*; *Abe H et al., 2008*; *Kawaoka S et al., 2011*). The monarch genome scaffold continuity (N50 = 0.207 Mb versus N50 = 14.2 Mb for *T. ni*; [*Zhan S et al., 2011*]) is insufficient to permit assembly of a W chromosome. Our genome assembly includes the 2.92 Mb *T. ni* W chromosome comprising 32 contigs (contig N50=101 kb). In *T. ni*, W-linked contigs have higher repeat content, lower gene density, and lower transcriptional activity than autosomal or Z-linked contigs (Figure 2B). Other lepidopteran W chromosomes are similarly enriched in repeats and depleted of genes (*Abe H et al., 2005*; *Fuková I et al., 2005*; *Traut W et al., 2007*).

A search for *T. ni* genes that are homologous to insect sex determination pathway genes detected *doublesex* (*dsx*), *masculinizer* (*masc*), *vitellogenin, transformer 2, intersex, sex lethal, ovarian tumor, ovo*, and *sans fille*. *T. ni* males produce a four-exon isoform of *dsx*, while females generate a six-exon *dsx* isoform (Figure 2–figure supplement 1D). The Lepidoptera-specific gene *masc* encodes a CCCH zinc finger protein. *masc* is associated with the expression of the sex-specific isoforms of *dsx* in lepidopterans, including silkworm (*Katsuma S et al., 2015*). As in *B. mori, T. ni masc* lies next to the *scap* gene, supporting our annotation of *T. ni masc*. Lepidopteran *masc* genes are rapidly diverging and have low (30.1%) sequence identity with one another. Figure 2C shows the multiple sequence alignment of the CCCH zinc finger domain of Masc proteins from several lepidopteran species.

### Telomeres and centromeres

Like many non-dipteran insects, *T. ni* has a single telomerase gene and telomeres containing TTAGG repeats (*Sahara K et al., 1999*). We found 40 (TTAGG)n stretches longer than 100 nt (mean ± S.D. = 600 ± 800 nt), nine at and 31 near contig boundaries (Supplementary file 1F; distance = 5,000 ± 6,000 nt for the 40 stretches), indicating that our assembly captures the sequences of many telomeres. More than half (59%) of the sequences flanking the (TTAGG)n repeats are transposons, and ~49% of these transposons belong to the non-long-terminal-repeat LINE/R1 family (Supplementary file 1G). These telomeric and subtelomeric characteristics of *T. ni* resemble those of *B. mori* (*Fujiwara H et al., 2005*).

Lepidopteran chromosomes generally lack a coherent, monocentric centromere and are instead holocentric or diffuse (*Labbé R et al., 2011*), and the silkworm, monarch butterfly, and diamondback moth genomes do not encode CenH3, a protein associated with monocentric chromosomes. The *T. ni* genome similarly does not contain a gene for CenH3, suggesting that its chromosomes are also holocentric.

### CpG content and DNA methylation

The *T. ni* genome is 35.6% GC, slightly less than *B. mori* (37.3%). The distributions of observed/expected CpG ratios in genes and across the genome (Figure 2— supplement 2A) reveal that *T. ni* is similar to other lepidopterans (silkworm, monarch butterfly, the diamondback moth) and a coleopteran species (the red flour beetle, *T*. *castaneum*), but different from honeybee and fruit fly. The honeybee genome has a high CpG content in genes and exhibits a bimodal CpG distribution across the genome as a whole; the fruit fly genome is uniformly depleted of CpG dinucleotides. The differences in CpG patterns reflect the presence of both the DNMT1 and DNMT3 DNA methyltransferases in the honeybee, the absence of either in fruit fly, and the presence of only DNMT1 in *T. ni, B. mori, D. plexippus, P. xylostella*, and *T. castaneum*. Thus, like many other insects, the *T. ni* genome likely has low levels of DNA methylation (*Xiang H et al., 2010*; *Glastad KM et al., 2011*).

### Transposons and repeats

The *T. ni* genome contains 75.3 Mb of identifiable repeat elements (20.5% of the assembly), covering 458 repeat families (Figure 2–figure supplement 2B, Supplementary file 1H). With this level of repeat content, *T. ni* fits well with the positive correlation between genome size and repeat content among lepidopteran genomes (Figure 2–figure supplement 2C).

The DNA transposon piggyBac was originally isolated from a *T. ni* cell line (*Fraser MJ et al., 1983*) and transposes effectively in a variety of species (*Lobo N et al., 1999*; *Bonin CP* and *Mann RS, 2004*; *Wang W et al., 2008*). We identified 262 copies of piggyBac in the Hi5 cell genome assembly. The family divergence rate of piggyBac is ~0.17%, substantially lower than other transposon families in the genome (Supplementary file 1I provides divergence rates for all transposon families). Among the individual piggyBac elements in the *T. ni* genome, 71 are specific to Hi5 cells. Compared to the 191 piggyBac insertions shared between *T. ni* and Hi5 cells (divergence rate = 0.22%), the Hi5 cell-specific elements are more highly conserved (divergence rate = 0.04%). We conclude that the piggyBac transposon entered the *T. ni* genome more recently than other transposons and, likely driven by the presence of many active piggyBac elements, expanded further during the immortalization of Hi5 cells in culture.

### microRNAs

microRNAs (miRNAs) are ~22 nt non-coding RNAs that regulate mRNA stability and translation (*He L* and *Hannon GJ, 2004*; *Gao G et al., 2005*). In insects, miRNA targets function in metamorphosis, reproduction, diapause, and other pathways of insect physiology and development (*Lucas K* and *Raikhel AS, 2013*). To characterize the *T. ni* miRNA pathway, we manually curated miRNA biogenesis genes such as *dcr-1, pasha, drosha*, and *ago2* (Supplementary file 2A) in testis, ovary, thorax, and Hi5 cells. To identify and annotate *T. ni* miRNAs, we sequenced small RNA from ovary, testis, thorax, and Hi5 cells. We predicted 295 miRNA genes (Figure 3, Supplementary file 3A and Supplementary file 4), including 77 conserved, 31 Lepidoptera-specific, and 187 novel, *T. ni*-specific miRNAs.

**Figure 3.**
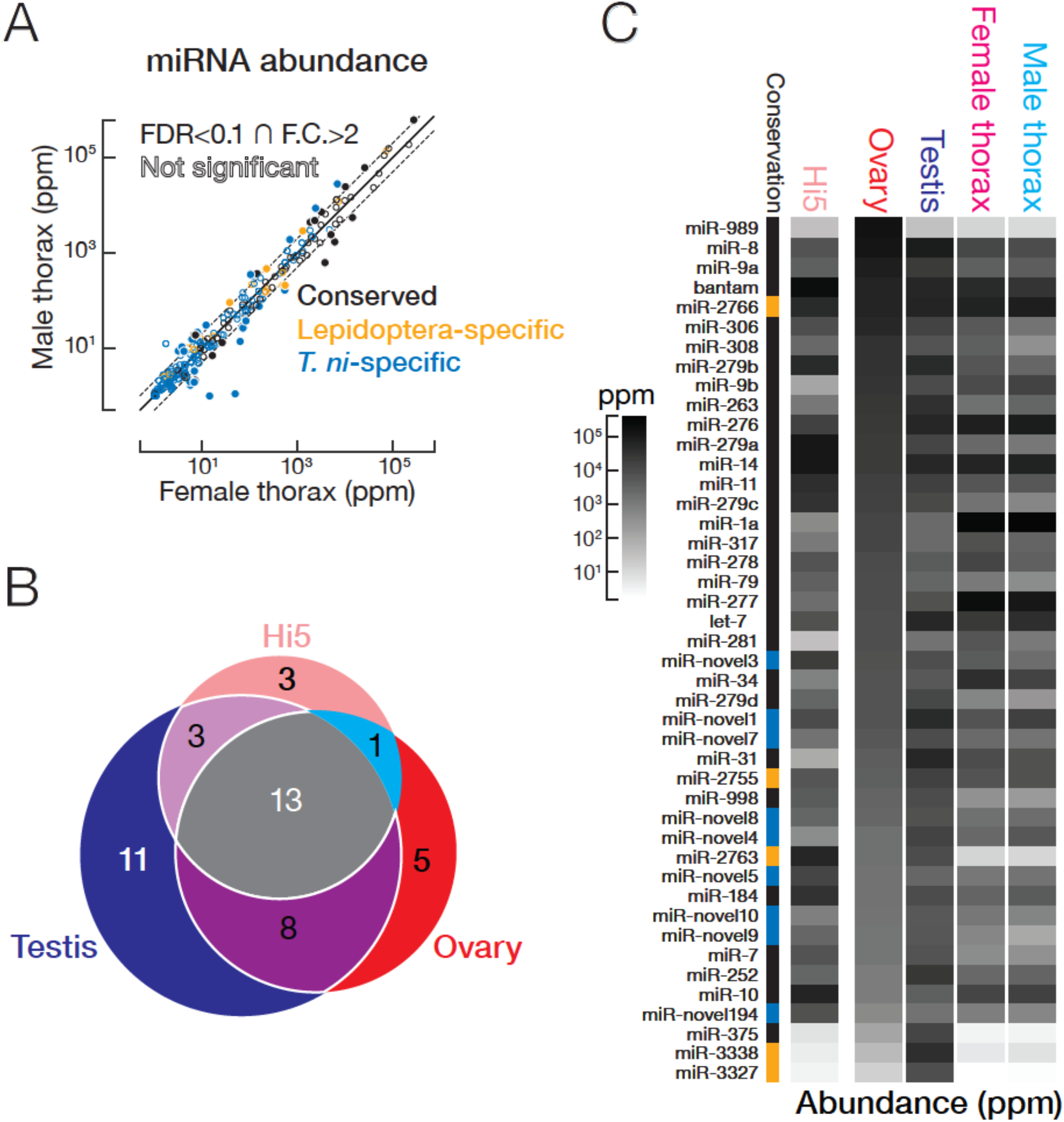
miRNA expression in *T. ni*. (A) Comparison of miRNA abundance in male and female *T. ni* thoraces. Solid circles, miRNAs with FDR < 0.1 and fold change > 2. Outlined circles, all other miRNAs. (B) Comparison of the tissue distribution of the 44 most abundant miRNAs among *T. ni* ovaries, testes, and Hi5. (C) Heat map showing the abundance of miRNAs in (B). miRNAs are ordered according to abundance in ovary. Conservation status uses the same color scheme in (A).

In thorax, 222 of 270 miRNAs had comparable abundance in males and females (≤2-fold difference and/or false discovery rate [FDR] ≥0.1; Figure 3A). Of the 48 miRNAs having significantly different abundances in female and male thorax (>2-fold difference and FDR <0.1; Figure 3A), miR-1a, let-7, and miR-278 were highly abundant (>1000 parts per million [ppm]) in either female or male thorax. miR-1a, a miRNA thought to be expressed in all animal muscle, was the most abundant miRNA in thorax in both sexes, but is 2.2-fold more abundant in males. miR-1 was previously shown to regulate muscle development in fruit flies (*Sokol NS* and *Ambros V, 2005*) and to increase when locusts transition from solitary to swarming (*Wei Y et al., 2009*). *T. ni* let-7, which has the same mature miRNA sequence as its *D. melanogaster, C. elegans*, and mammalian counterparts (*Lagos-Quintana M et al., 2001*) was also more abundant in males, whereas miR-278 was 2.6-fold more abundant in females. let-7 may act in sex-specific pathways in metamorphosis (*Caygill EE* and *Johnston LA, 2008*), whereas miR-278 may play a sex-specific role in regulating energy homeostasis (*Teleman AA et al., 2006*).

A subset of less well conserved miRNAs was also differentially expressed between male and female thorax. In general, poorly conserved miRNAs were less abundant: the median expression level for conserved miRNAs was 316 ppm, but only 161 ppm for Lepidoptera-specific and 4.22 ppm for *T. ni*-specific miRNAs. However, mir-2767, a Lepidoptera-specific miRNA, and three *T. ni*-specific miRNAs (mir-novel1, mir-novel4, mir-novel11) were both abundant (>1000 ppm) and differentially expressed in males and female thorax. We speculate that these recently evolved miRNAs may prove useful as targets for pest management.

Ovary, testis, and Hi5 cells have distinct miRNA expression profiles. We analyzed the expression patterns of the 44 most abundant miRNAs (Figure 3B and 3C), which explain 90% of miRNA reads in a tissue or cell line. Thirteen were expressed in ovaries, testes, and Hi5. Of these 13, 11 were significantly more abundant in testis, 5 in ovary, and 3 in Hi5 cells (Figure 3B), suggesting that these miRNAs have important tissue-or cell-type-specific roles. miR-31 and miR-375, highly expressed in *T. ni* testis, are both mammalian tumor suppressors (*Creighton CJ et al., 2010*; *Kinoshita T et al., 2012*). miR-989, the most abundant miRNA in *T. ni* ovaries, plays an important role in border cell migration during *Drosophila* oogenesis (*Kugler J-M et al., 2013*). miR-10, a miRNA in the Hox gene cluster, was preferentially expressed in Hi5 cells; its orthologs have been implicated in development and cancer (*Lund AH, 2009*), suggesting miR-10 played a role in the immortalization of the germline cells from which Hi5 cells derive.

### siRNAs

siRNAs, typically 20–22 nt long, regulate gene expression, defend against viral infection, and silence transposons (*Sanchez-Vargas I et al., 2009*; *van Rij RP et al., 2006*; *Chung W-J et al., 2008*; *Tyler DM et al., 2008*; *Zambon RA et al., 2006*; *Okamura K et al., 2008b*; *Okamura K et al., 2008b*; *Flynt A et al., 2009*). They are processed by Dicer from double-stranded RNAs or hairpins into short double-stranded fragments bearing two-nucleotide, overhanging 3′ ends, which are subsequently loaded into Argonaute proteins (*Bernstein E et al., 2001*; *Elbashir SM et al., 2001*; *Siomi H and Siomi MC, 2009*). siRNAs direct Argonaute proteins to cleave their targets via extensive sequence complementarity.

### Endogenous siRNAs from transposons and *cis*-NATs

Endogenous siRNAs (endo-siRNAs) can derive from transposon RNAs, *cis*-natural antisense transcripts (*cis*-NATs), and long hairpin RNAs (*Chung W-J et al., 2008*; *Czech B et al., 2008*; *Ghildiyal M et al., 2008*; *Okamura K et al., 2008a*; *Kawamura Y et al., 2008*; *Okamura K et al., 2008a*; *Tam OH et al., 2008*; *Watanabe T et al., 2008*) (hpRNAs). In *T. ni* ovary, testis, thorax, and Hi5 cells, 20.7–52.4% of siRNAs map to transposons, suggesting *T. ni* endogenous siRNAs suppress transposons in both the soma and the germline. Among the non-transposon siRNAs, <4.6% map to predicted hairpins, while 11.6–31.3% siRNAs map to *cis*-NATs (supplementary file 3B).

### Exogenous siRNAs against a virus

Hi5 cells are latently infected with a positive-sense, bipartite alphanodavirus, TNCL virus (*Li T-C et al., 2007*;) (Tn5 Cell Line virus). We asked if TNCL virus RNA is present in our *T. ni* samples and whether the RNAi pathway provides anti-viral defense via TNCL virus-derived siRNAs. We detected no viral RNA in the *T. ni* ovary, testis, or thorax transcriptome, but both TNCL virus RNA1 (5,010 fragments per kilobase of transcript per million mapped reads [FPKM]) and RNA2 (8,280 FPKM) were readily found in the Hi5 transcriptome (Figure 4A). To test whether Hi5 cells mount an RNAi defense to TNCL virus infection, we mapped small RNA-seq reads that were not mappable to the *T. ni* genome to the two TNCL virus genomic segments. TNCL virus mapping small RNAs showed a median length of 21 nt (modal length = 20 nt; Figure 4A), typical for siRNAs, suggesting that the Hi5 RNAi pathway actively combats the virus. The TNCL virus-mapping small RNAs bear the two-nucleotide, 3′ overhanging ends that are the hallmark of siRNAs (Figure 4B; [*Elbashir SM et al., 2001*; *Elbashir SM et al., 2001*; *Elbashir SM et al., 2001*]. Moreover, the phased pattern of TNCL virus-mapping siRNAs suggests they are made one-after-another starting at the end of a dsRNA molecule: the distance between siRNA 5′ ends shows a periodicity of 20 nt, the length of a typical TNCL virus-mapping siRNA (Figure 4C). In *D. melanogaster*, Dicer-2 processively produces siRNAs, using ATP energy to translocate along a dsRNA molecule (*Cenik ES et al., 2011*). The phasing of anti-viral siRNAs in Hi5 cells suggests that *T. ni* Dicer-2 similarly generates multiple siRNAs from each molecule of dsRNA before dissociating.

**Figure 4.**
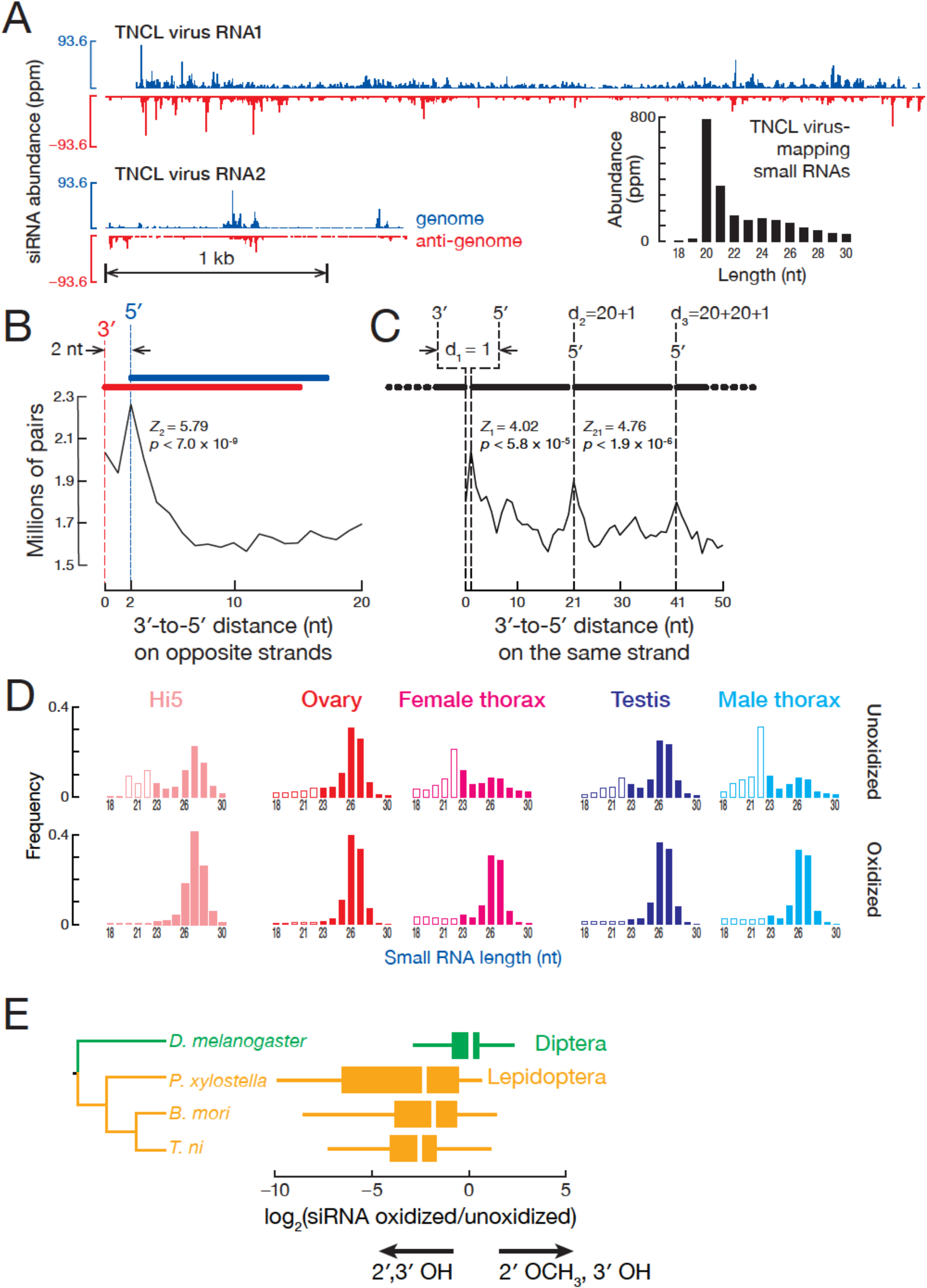
siRNA. (A) Distribution of siRNAs mapping to TNCL virus in the genomic (blue) and anti-genomic orientation (red). Inset: length distribution of TNCL virus-mapping small RNAs. (B) Distance between the 3′ and 5′ ends of siRNAs on opposite viral strands. (C) Distance between the 3′ and 5′ ends of siRNAs on the same viral strand. (D) Length distribution of small RNAs from unoxidized and oxidized small RNA-seq libraries. (E) Lepidopteran siRNAs are not 2′-*O*-methylated. The box plots display the ratio of abundance (as a fraction of all small RNAs sequenced) for each siRNA in oxidized versus unoxidized small RNA-seq libraries. The tree shows the phylogenetic relationships of the analyzed insects. Outliers are not shown.

### Lepidopteran siRNAs are not 2′-*O*-methylated

The discovery that the 3′ ends of *D. melanogaster* siRNAs, but not miRNAs, are 2′-*O*-methylated (*Pelisson A et al., 2007*) led to the idea that insects in general methylate both siRNAs and piRNAs. Resistance to oxidation by NaIO_4_ is the hallmark of 3′ terminal, 2′-*O*-methylation, and the enrichment of a small RNA in a high-throughput sequencing library prepared from NaIO4-treated RNA suggests 2′-*O*-methylation. Conversely, depletion of small RNAs, such as miRNAs, from such an oxidized RNA library is strong evidence for unmodified 2′,3′ vicinal hydroxyl groups. Surprisingly, TNCL virus-mapping siRNAs were 130-fold depleted from our oxidized small RNA-seq library (22.0 ppm) compared to the unoxidized library (2,870 ppm), suggesting that they are unmethylated. Sequencing of oxidized and unoxidized small RNA from *T. ni* ovary, testis, and thorax detected 20–22 nt peaks in unoxidized libraries; such peaks were absent from oxidized libraries (Figure 4D), suggesting that *T. ni* genome-mapping, endogenous siRNAs also lack 2′-O-methylation. We conclude that both *T. ni* exo-and endo-siRNAs are not 2′-*O*-methyl modified.

Are siRNAs unmethylated in other Lepidopteran species? We sequenced oxidized and unoxidized small RNAs from two additional Lepidoptera: *P. xylostella* and *B. mori*. Like *T. ni*, siRNAs from these Lepidoptera were abundant in libraries prepared from unoxidized small RNA but depleted from oxidized libraries (Figure 4–figure supplement 1A). The ratio of siRNAs in the oxidized library to siRNAs in the corresponding unoxidized library (ox/unox) provides a measure of siRNA 2′,3′ modification. For *D. melanogaster* siRNAs, the median ox/unox ratio was 1.00, whereas the three Lepidoptera species had median ox/unox ratios between 0.17 and 0.22 (Figure 4E), indicating their siRNAs were depleted from oxidized libraries and therefore bear unmodified 2′,3′ hydroxyl groups. We conclude that the last common ancestor of *T. ni, B. mori*, and *P. xylostella,* which diverged 170 mya, lacked the ability to 2′-*O*-methylate siRNA 3′ ends. We do not currently know whether the last common ancestor of Lepidoptera lost the capacity to methylate siRNAs or if some or all members of Diptera, the sister order of Lepidoptera, acquired this function, which is catalyzed by the piRNA-methylating enzyme Hen1 (*Saito K et al., 2007*; *Horwich MD et al., 2007*; *Kirino Y* and *Mourelatos Z, 2007*).

Terminal 2′ methylation of *D. melanogaster* siRNAs is thought to protect them from non-templated nucleotide addition (tailing), trimming, and wholesale degradation (*Ameres SL et al., 2010*). Since *T. ni* siRNAs lack a 2′-*O*-methyl group at their 3′ ends, we first asked if we could observe frequent trimming by examining shorter TNCL-mapping siRNA (18-19 nt). These siRNAs could account for 1.05% of all TNCL-mapping siRNAs. They did not possess the one-after-another pattern (Z1 = -0.674, *p* = 0.500), yet more than 97.5% of these were prefixes of longer, phased siRNAs, indicating that these were trimmed siRNAs. We conclude that TNCL siRNA trimming is rare in Hi5 cells. We next asked whether *T. ni* and other lepidopteran siRNAs have higher frequencies of tailing. Despite the lack of 2′-*O*-methylation, most TNCL virus siRNAs were not tailed: just 6.69% of all virus-mapping small RNA reads contained 3′ non-templated nucleotides (Figure 4–figure supplement 1B). Among the 3′ non-templated nucleotides, the most frequent addition was one or more uridines (49.6%) as observed previously for miRNAs and siRNAs in other species (*Ameres SL et al., 2010*; *Chou MT et al., 2015*). Endogenous siRNA tailing frequencies for the lepidopterans *T. ni* (10.2%, ovary), *B. mori* (5.97%, eggs), and *P. xylostella* (8.58%, ovary) were also similar to *D. melanogaster* (6.71%, ovary). We speculate that lepidopterans have other mechanisms to maintain siRNA stability or that trimming and tailing in lepidopterans are less efficient than in flies.

### piRNAs

In animals, piRNAs, ~23–32 nt long, protect the germline genome by suppressing the transcription or accumulation of transposon and repetitive RNA (*Girard A et al., 2006*; *Lau NC et al., 2006*; *Vagin VV et al., 2006*; *Aravin AA et al., 2007*; *Brennecke J et al., 2007*). In *D. melanogaster*, dedicated transposon-rich loci (piRNA clusters) give rise to piRNA precursor transcripts, which are processed into piRNAs that are then loaded into one of three PIWI proteins, Piwi, Aubergine (Aub), or Argonaute3 (Ago3). Piwi acts in the nucleus to direct tri-methylation of histone H3 on lysine 9 of H3K9me3 on transposons and repetitive genomic DNA (*Le Thomas A et al., 2014*; *Le Thomas A et al., 2014*). In fly cytoplasm, piRNAs guide the Piwi paralog Aub to cleave transposon mRNAs. The mRNA cleavage products can then produce more piRNAs, which are loaded into Ago3. In turn, these sense piRNAs direct Ago3 to cleave transcripts from piRNA clusters, generating additional piRNAs bound to Aub. The resulting “Ping-Pong” feed-forward loop both amplifies piRNAs and represses transposon activity (*Brennecke J et al., 2007*; *Gunawardane LS et al., 2007*). Finally, Ago3 cleavage not only produces Aub-bound piRNAs, but also initiates the production of Piwi-bound, phased piRNAs that diversify the piRNA pool (*Han BW et al., 2015a*; *Mohn F et al., 2015*).

### piRNA pathway proteins

The *T. ni* genome contains a full repertoire of genes encoding piRNA pathway proteins (Supplementary file 2B). These genes were expressed at higher levels in ovary, testis, and Hi5 cells compared to somatic tissues (median ratios: ovary/thorax = 14.2, testis/thorax = 2.9, and Hi5/thorax = 4.9; Figure 5A), suggesting that piRNAs function in the *T. ni* germline and that the ovary-derived Hi5 cell line recapitulates the germline piRNA pathway. Although most *T. ni* piRNA pathway genes correspond directly to their *D. melanogaster* orthologs, *T. ni* encodes only two PIWI proteins, Ciwi (i.e., TnPiwi) and TnAgo3. The fly proteins Aub and Piwi are paralogs that arose from a single ancestral PIWI protein after the divergence of flies and mosquitos (*Lewis SH et al., 2016*). We do not yet know whether Ciwi functions more like *Drosophila* Aub or Piwi. In *D*. *melanogaster*, piRNA clusters—the genomic sources of most transposon-silencing germline piRNAs—are marked by the proteins Rhino, Cutoff, and Deadlock, which allow transcription of these heterochromatic loci (*Klattenhoff C et al., 2009*; *Zhang Z et al., 2014*). *T. ni* lacks detectable Rhino, Cutoff, and Deadlock orthologs. In fact, this trio of proteins is poorly conserved, and the mechanism by which they mark fly piRNA source loci may be unique to Drosophilids. In this regard, *T. ni* likely provides a more universal insect model for the mechanisms by which germ cells distinguish piRNA precursor RNAs, which are processed into mature piRNAs, from other protein-coding and non-coding transcripts.

**Figure 5.**
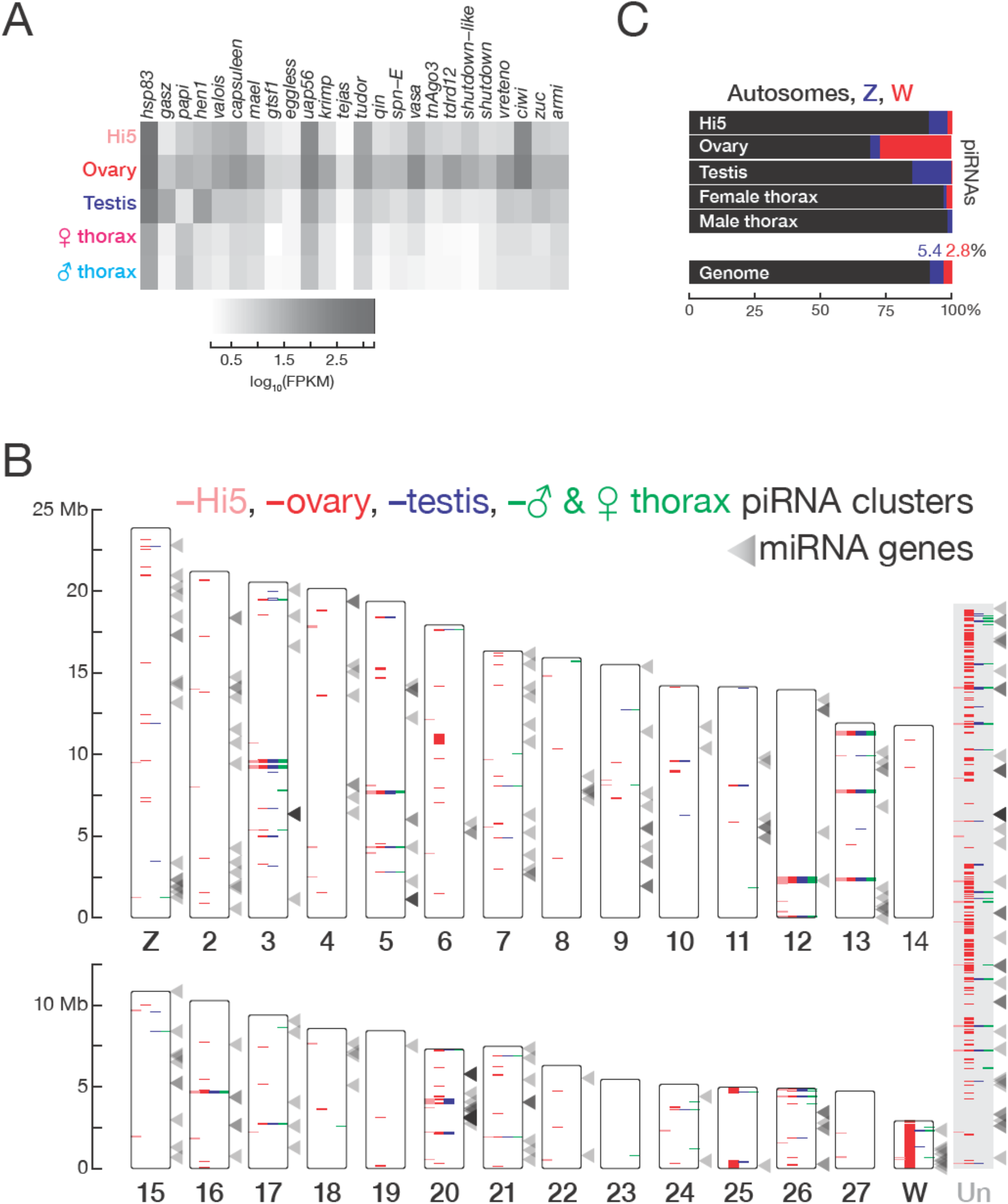
piRNAs and miRNAs in the *T. ni* genome. (A) Abundance of mRNAs encoding piRNA pathway proteins) in Hi5 cells, ovary, testis, and thorax. (B) Ideogram displaying the positions of miRNA genes (arrowheads) and piRNA clusters in the *T. ni* genome. Color-coding reports tissue expression for Hi5 cells, ovaries, testis, and thorax. Contigs that cannot be placed onto chromosome-length scaffolds are arbitrarily concatenated and are marked ‘Un.’ (C) Distribution of piRNAs among the autosomes, Z, and W chromosomes in Hi5 cells, ovary, testis, and female and male thorax, compared with the fraction of the genome corresponding to autosomes, W, and Z chromosomes.

### piRNA cluster architecture

In both the germline and the soma, *T. ni* piRNAs originate from discrete genomic loci. To define these piRNA source loci, we employed an expectation-maximization algorithm that resolves piRNAs mapping to multiple genomic locations. Applying this method to multiple small RNA-seq datasets, we defined piRNA-producing loci comprising 10.7 Mb (332 clusters) in ovary, 3.1 Mb (76 clusters) in testis, 3.0 Mb (71 clusters) in Hi5 cells, and 2.4 Mb (65 clusters) in thorax (Figure 5B). For each tissue or cell-type, these 393 clusters explain >70% of uniquely mapped piRNAs and >70% of all piRNAs when using expectation-maximization mapping. A core set of piRNA-producing loci comprising 1.5 Mb is active in both germline and somatic tissues.

*T. ni* piRNA clusters vary substantially in size and expression level. In ovary, half the bases in piRNA clusters are in just 67 loci, with a median length of 53 kb. Among these, five span >200 kb, while the smallest is just 38 kb. The most productive piRNA source is a 264 kb locus on chromosome 13 (Figure 5–figure supplement 1); 7.8% of uniquely mapped piRNAs—50,000 distinct piRNA sequences—reside in this locus. Collectively, the top 20 ovary piRNA loci explain half the uniquely mapped piRNAs, yet constitute only 0.7% of the genome. Globally, 61.9% of bases in piRNA clusters are repetitive, and 74.5% transposon-mapping piRNAs are antisense, suggesting that *T. ni* uses antisense piRNAs to suppress transposon transcripts.

In the fly ovary germline, most piRNA clusters generate precursor RNAs from both DNA strands. These dual-strand clusters fuel the ‘Ping-Pong’ amplification cycle (*Brennecke J et al., 2007*; *Gunawardane LS et al., 2007*). Other fly piRNA clusters, such as the paradigmatic *flamenco* gene (*Prud’homme N et al., 1995*; *Brennecke J et al., 2007*; *Pelisson A et al., 2007*; *Malone CD et al., 2009*; *Goriaux C et al., 2014*) are transcribed from one strand only and are organized to generate antisense piRNAs directly, without further Ping-Pong amplification (*Malone CD et al., 2009*). These uni strand clusters are the only sources of piRNAs in the follicle cells, somatic cells that support fly oocyte development and express only a single PIWI protein, Piwi (*Malone CD et al., 2009*).

The *T. ni* genome contains both dual- and uni-strand piRNA clusters. In ovary, 67 of 332 piRNA-producing loci are dual-strand (Watson/Crick >0.5 and Watson/Crick < 2). These loci produce 36.8% of uniquely mapped piRNAs and 23.4% of all piRNAs; 71.7% of transposon-mapping piRNAs from these loci are antisense. The remaining 265 uni-strand loci account for 54.0% of uniquely mapped piRNAs and 36.1% of all piRNAs. Most piRNAs (74.8%) from uni-strand clusters are antisense to transposons, the orientation required for repressing transposon mRNA accumulation. At least part of the piRNA antisense bias reflects positive selection for antisense insertions in unistrand clusters: 57.2% of transposons are inserted opposite to the direction of piRNAs precursor transcription (Figure 5–figure supplement 2A). For example, 99.0% of piRNAs that can be uniquely assigned to a 77 kb uni-strand cluster on chromosome 20 are from the Crick strand, while 79.7% of transposon sequences at this locus lie on the Watson strand.

### Nearly the entire W chromosome produces piRNAs

The largest ovary cluster is a 462 kb W-linked region, consistent with our finding that the W chromosome is a major source of piRNAs (Figure 5B and 5C and Figure 5— figure supplement 2B). Our data likely underestimates the length of this large piRNA cluster, as it is difficult to resolve reads mapping to its flanking regions: 70.8% of bases in the flanking regions do not permit piRNAs to map uniquely to the genome. In fact, 85.1% of the sequences between clusters on the W chromosome are not uniquely mappable. These gaps appear to reflect low mappability and not boundaries between discrete clusters. Instead, we propose that the entire W chromosome is a giant piRNA cluster.

To further test this idea, we identified piRNA reads that uniquely map to one location among all contigs and measured their abundance per kilobase of the genome. W-linked contigs had a median piRNA abundance of 14.4 RPKM in ovaries, 379-fold higher than the median of all autosomal and Z-linked contigs, consistent with the view that almost the entire W chromosome produces piRNAs. In *B. mori* females, a plurality of piRNAs come from the W chromosome: ovary-enriched piRNAs often map to W-linked BAC sequences, but not autosomes (*Kawaoka S et al., 2011*). Similarly, for *T. ni*, 27.2% of uniquely mapping ovary piRNAs derive from W-linked sequences, even though these contigs compose only 2.8% of the genome (Figure 5C). The W chromosome may produce more piRNAs than our estimate, as the unassembled repetitive portions of the W chromosome likely also produce piRNAs. Thus, the entire W chromosome is a major source of piRNAs in *T. ni* ovaries (Figure 5B). To our knowledge, the *T. ni* W chromosome is the first example of an entire chromosome devoted to piRNA production.

In contrast to the W chromosome, *T. ni* autosomes and the Z chromosome produce piRNAs from discrete loci—63 autosomal and 11 Z-linked contigs had piRNA levels >10 rpkm. Few piRNAs are produced outside of these loci: for example, the median piRNA level across all autosomal and Z-linked contigs was ~0 in ovaries (Figure 5–figure supplement 2B).

### Expression of piRNA clusters

In the *T. ni* germline, piRNA production from individual clusters varies widely, but the same five piRNA clusters produce the most piRNAs in ovary (34.9% of piRNAs), testis (49.3%), and Hi5 cells (44.0%), suggesting that they serve as master loci for germline transposon silencing. Other piRNA clusters show tissue-specific expression, with clusters on the W chromosome (probably as one single giant cluster) producing more piRNAs in ovary than in Hi5 cells, and three Z-linked clusters producing many more piRNAs in testis than in ovary (15.0–24.7 times more), even after accounting for the absence of dosage compensation in germline tissues (Figure 6–figure supplement 1A).

**Figure 6.**
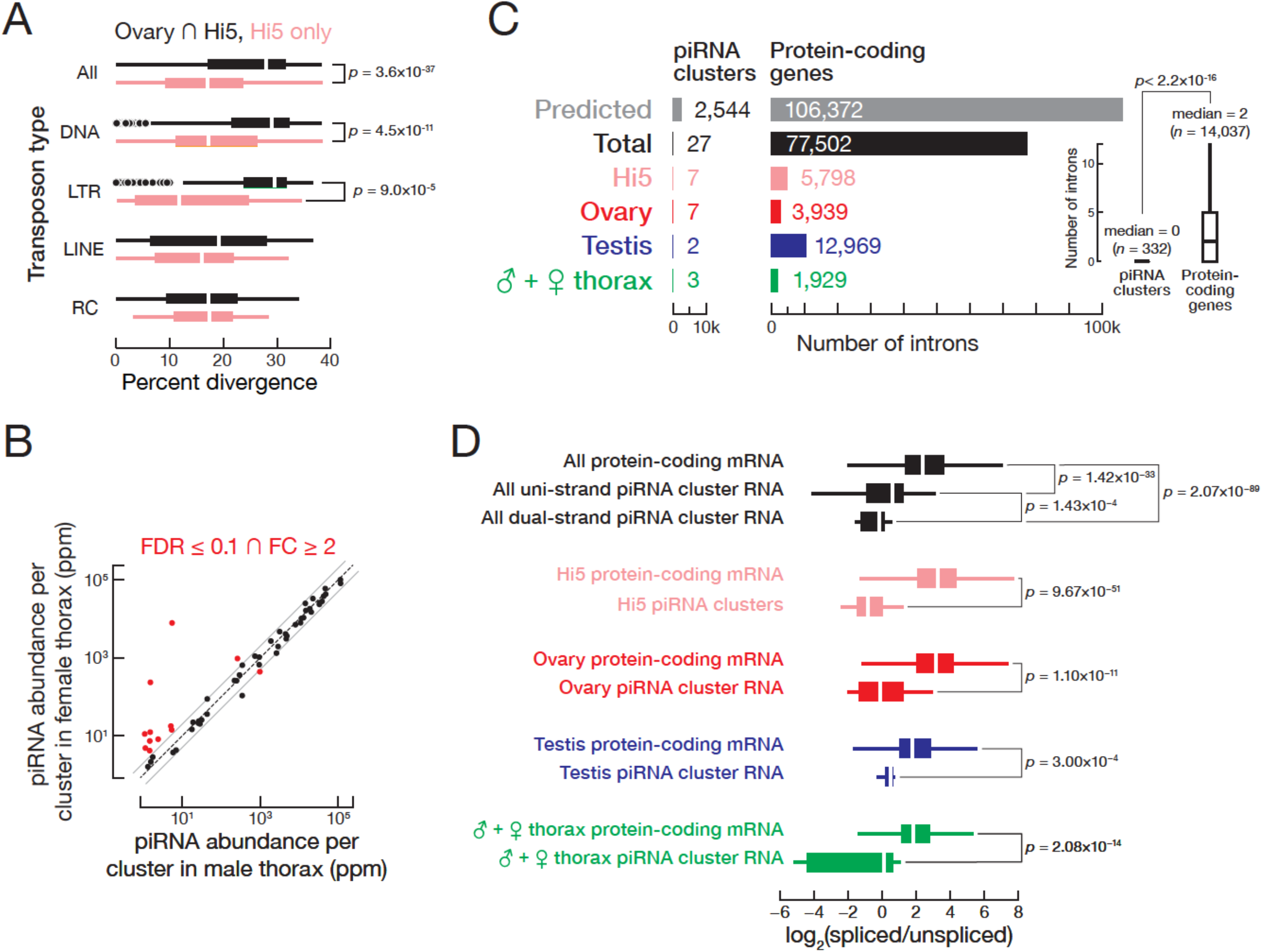
(A) Hi5-specific piRNA clusters contain younger transposon copies. RC, rolling-circle transposons; LINE, Long interspersed nuclear elements; LTR, long terminal repeat retrotransposon; DNA, DNA transposon. (B) Comparison of piRNA abundance per cluster in female and male thorax. (C) piRNA precursors are rarely spliced. The number of introns supported by exon-exon junction-mapping reads is shown for protein-coding genes and for piRNA clusters for each tissue or cell type. (D) piRNA precursors are inefficiently spliced. Splicing efficiency is defined as the ratio of spliced over unspliced reads. Splice sites were categorized into those inside and outside piRNA clusters. Outlier are not shown.

Hi5 cells are female, yet many piRNA-producing regions of the W chromosome that are active in the ovary produce few piRNAs in Hi5 cells (Figure 6–figure supplement 1A). We do not know whether this reflects a reorganization of cluster expression upon Hi5 cell immortalization (*Granados RR et al., 1986*) or if Hi5 cluster expression corresponds to a specific germ cell type that is underrepresented in whole ovaries. At least 40 loci produce piRNAs in Hi5 cells but not the *T. ni* ovary. Comparison of DNA-seq data from *T. ni* and Hi5 identified 74 transposon insertions in 12 of the Hi5-specific piRNA clusters. Older transposons have more time to undergo sequence drift from the consensus sequence of the corresponding transposon family. The 74 Hi5-specific transposon insertions, which include both DNA and LTR transposons, had significantly lower divergence rates than those common to ovary and Hi5 cells (Figure 6A), consistent with the idea that recent transposition events generated the novel piRNA clusters in Hi5 cells. We conclude that the Hi5-specific piRNA-producing loci are quite young, suggesting that *T. ni* and perhaps other lepidopterans can readily generate novel piRNA clusters.

piRNA clusters active in thorax occupy ~0.57% of the genome and explain 86.8% of uniquely mapped somatic piRNAs in females and 89.5% in males. More than 90% of bases in clusters expressed in thorax are shared with clusters expressed in ovary (Figure 6–figure supplement 1B), and such broadly expressed clusters explain 83.7% of uniquely mapping piRNAs in female thorax and 86.1% in male thorax. Thus, the majority of piRNAs in the *T. ni* soma come from clusters that are also active in the germline. Autosomal piRNA cluster expression is similar between female and male thorax; among the 12 clusters differentially expressed between male and female thorax, nine W-linked clusters produce significantly more piRNAs in female thorax (Figure 6B).

### piRNA precursor transcripts are rarely spliced

In *D. melanogaster*, Rhino suppresses splicing of piRNA precursors transcribed from dual-strand piRNA clusters (*Mohn F et al., 2014*; *Zhang Z et al., 2014*). Fly uni-strand piRNA clusters do not bind Rhino and behave like canonical RNA polymerase II transcribed genes (*Brennecke J et al., 2007*; *Goriaux C et al., 2014*). Although *T. ni* has no *rhino* ortholog, its piRNA precursor RNAs are rarely spliced as observed in flies. We identified splicing events using RNA-seq reads aligned to the genome in a split manner, requiring ≥10 reads that map across exon-exon junctions and a minimum splicing entropy of 2 to exclude PCR duplicates (*Graveley BR et al., 2011*). This approach detected just 27 splice sites among all piRNA precursor transcripts from ovary, testis, thorax, and Hi5 piRNA clusters (Figure 6C). Of these 27 splice sites, 19 fall in unistrand piRNA clusters. We conclude that, as in flies, transcripts from *T. ni* dual-strand piRNAs clusters are rarely if ever spliced. Unlike flies (*Goriaux C et al., 2014*), RNA from *T. ni* uni-strand piRNA clusters undergoes splicing infrequently.

The absence of piRNA precursor splicing in dual-strand piRNA clusters could reflect an active suppression of the splicing machinery or a lack of splice sites. To distinguish between these two mechanisms, we predicted gene models for piRNA-producing loci, employing the same parameters used for protein-coding gene prediction. For piRNA clusters, this approach generated 1,332 gene models encoding polypeptides >200 amino acids. These gene models comprise 2,544 introns with consensus splicing signals (Figure 6–figure supplement 1C). Notably, ~90% of these predicted gene models had high sequence similarity to transposon consensus sequences (blast e-value<10^‑10^), indicating that many transposons in piRNA clusters have intact splice sites. We conclude that piRNA precursors contain splice sites, but their use is actively suppressed.

To measure splicing efficiency, we calculated the ratio of spliced to unspliced reads for each predicted splice site in the piRNA clusters. High-confidence splice sites in protein-coding genes outside piRNA clusters served as a control. Compared to the control set of genes, splicing efficiency in piRNA loci was 9.67-fold lower in ovary, 2.41-fold lower in testis, 3.23-fold lower in thorax, and 17.0-fold lower in Hi5 cells (Figure 6D), showing that *T. ni* piRNA precursor transcripts are rarely and inefficiently spliced. In flies, uni-strand piRNA cluster transcripts are spliced, whereas splicing is suppressed for dual-strand cluster transcripts. To test whether uni- and dual-strand piRNA cluster transcripts are differentially spliced in *T. ni*, we evaluated the experimentally supported splice sites from Hi5, ovary, testis, and thorax collectively. Dual-strand cluster transcripts had significantly lower splicing efficiency (1.71-fold lower) compared to uni-strand clusters (Figure 6D). Thus, *T. ni* suppresses splicing of dual-strand piRNA cluster transcripts by a mechanism distinct from the Rhino-dependent pathway in *D. melanogaster*. That this novel splicing suppression pathway is highly efficient in Hi5 cells should facilitate its molecular dissection.

### Genome-editing and single-cell cloning of Hi5 cells

The study of arthropod piRNAs has been limited both by a lack of suitable cultured cell models and by the dominance of *D. melanogaster* as a piRNA model for arthropods generally. Although Vasa-positive *D. melanogaster* ovarian cells have been isolated and cultured (*Niki Y et al., 2006*), no dipteran germ cell line is currently available. *D*. *melanogaster* somatic OSS, OSC and Kc167 cells produce piRNAs, but lack key features of the canonical germline pathway (*Lau NC et al., 2009*; *Saito K et al., 2009*; *Vrettos N et al., 2017*). In addition to Hi5 cells, lepidopteran cell lines from *Spodoptera frugiperda* (Sf9) and *B. mori* (BmN4) produce germline piRNAs (*Kawaoka S et al., 2009*). The *S. frugiperda* genome remains a draft with 37,243 scaffolds and an N50 of 53.7 kb (*Kakumani PK et al., 2014*). Currently, the BmN4 cell line is the only ex vivo model for invertebrate germline piRNA biogenesis and function. The *B. mori* genome sequence currently comprises 43,463 scaffolds with an N50 of 4.01 Mb (*The ISG, 2008*). Unfortunately, BmN4 cells are difficult to grow and readily differentiate into two morphologically distinct cell types (*Iwanaga M et al., 2014*). Although genome editing with Cas9 has been demonstrated in BmN4 cells (*Zhu L et al., 2015*), no protocols for cloning individual, genome-modified BmN4 cells have been reported (*Mon H et al., 2004*; *Kawaoka S et al., 2009*; *Honda S et al., 2013*). In contrast, Hi5 cells cultured using commercially available media, readily transfected, and, we report here, efficiently engineered with Cas9 and grown from single cells into clonal cell lines.

The bacterial DNA nuclease Cas9, targeted by a single guide RNA (sgRNA) enables rapid and efficient genome editing in worms, flies, and mice, as well as in a variety of cultured animal cell lines (*Jinek M et al., 2012*; *Barrangou R* and *Horvath P, 2017*; *Komor AC et al., 2017*). The site-specific double-strand DNA breaks (DSBs) catalyzed by Cas9 can be repaired by error-prone non-homologous end joining (NHEJ), disrupting a protein-coding sequence or, when two sgRNAs are used, deleting a region of genomic DNA. Alternatively, homology-directed repair (HDR) using an exogenous DNA template allows the introduction of novel sequences, including fluorescent proteins or epitope tags, as well as specific point mutations in individual genes (*Cong L et al., 2013*).

As a proof-of-concept, we used Cas9 and two sgRNAs to generate an ~880 bp deletion in the piRNA pathway gene *ciwi*. The two sgRNAs, whose target sites lie 881 bp apart (Figure 7A), were transcribed *in vitro*, loaded into purified, recombinant Cas9 protein, and the resulting sgRNA/Cas9 ribonucleoprotein complexes (RNPs) were transfected into Hi5 cells. PCR of genomic DNA isolated 48 h later was used to detect alterations in the *ciwi* gene. A novel PCR product, ~900 bp smaller than the product amplified using DNA from untransfected Hi5 cells, indicated that the desired deletion had been created (Figure 7B). Sanger sequencing of the PCR products confirmed deletion of 881–896 bp from the *ciwi* gene. The presence of indels—short deletions and non-templated nucleotide additions—at the deletion junction is consistent with a Cas9-mediated dsDNA break having been repaired by NHEJ (Figure 7A).

**Figure 7.**
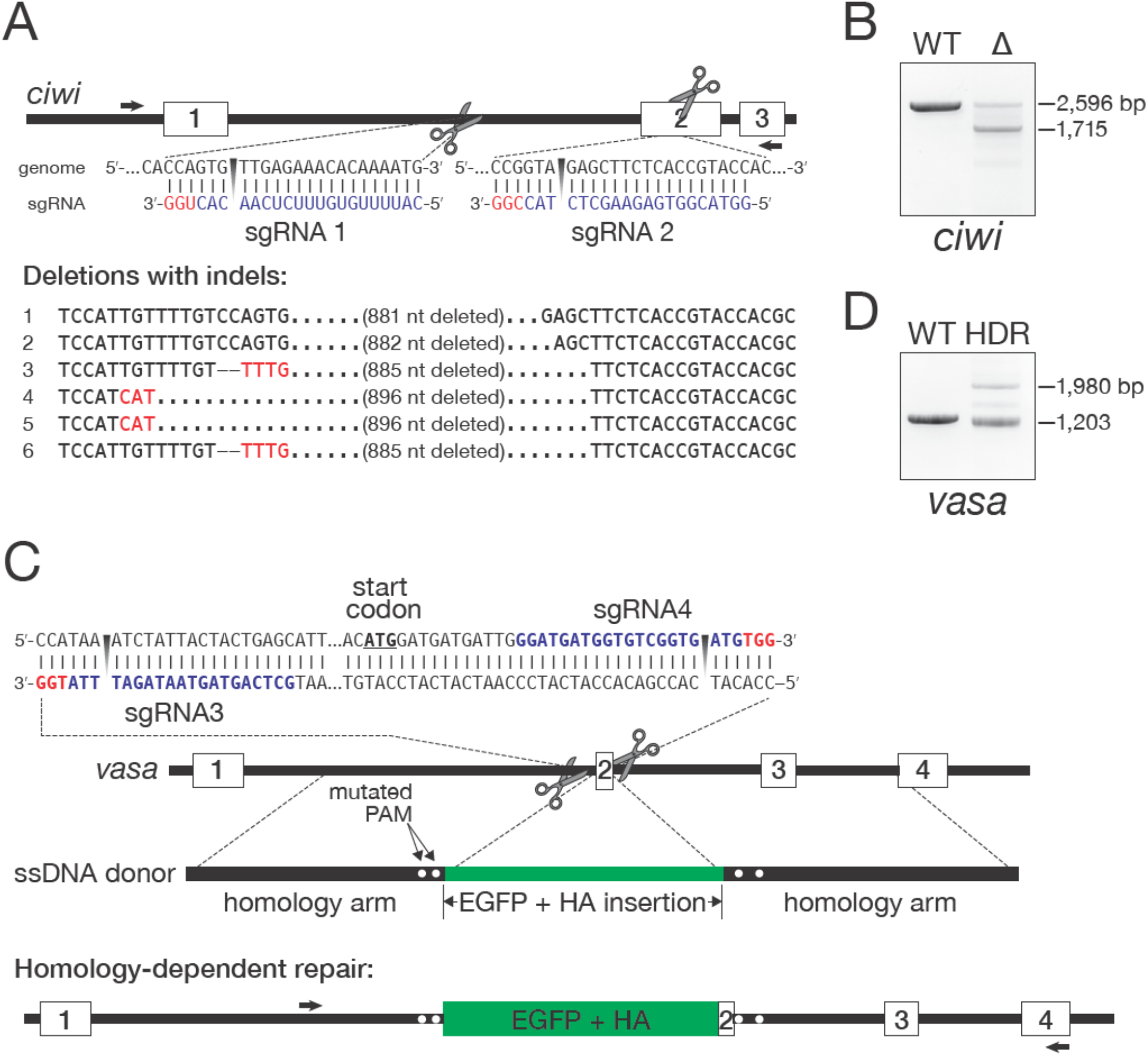
Genome editing in Hi5 cells. (A) Strategy for using Cas9/sgRNA RNPs to generate a loss-of-function *ciwi* deletion allele. Red, protospacer-adjacent motif (PAM); blue, protospacer sequence. Arrows indicate the diagnostic forward and reverse primers used in PCR to detect genomic deletions (Δ). Sanger sequencing of the ~1700 bp PCR products validated the *ciwi* deletions. (B) An example of PCR analysis of a *ciwi* deletion event. (C) Strategy for using Cas9/sgRNA RNPs and a single-stranded DNA homology donor to insert EGFP and an HA-tag in-frame with the *vasa* open reading frame. (D) An example of PCR analysis of a successful HDR event. DNA isolated from wild type (WT) and FACS-sorted, EGFP-expressing Hi5 cells (HDR) were used as templates.

As an alternative gene disruption strategy, we used an exogenous donor DNA to disrupt *ciwi* by inserting mCherry coding sequences 5ʹ to the *ciwi* start codon in exon 2. As for *vasa*, we designed two sgRNAs flanking the start codon (Figure 7–figure supplement 1A). As a donor, we used a plasmid DNA encoding mCherry and the *D*. *melanogaster p10* 3ʹ UTR (*Pfeiffer BD et al., 2012*). Cas9 and the two sgRNAs were cotransfected with the DNA donor, and, one week later, mCherry-positive cells were detected by fluorescence microscopy.

To establish a clonally derived line from the mCherry-disrupted *ciwi* Hi5 cells, individual mCherry-positive cells were isolated by FACS and cultured on selectively permeable filters above a feeder layer of wild-type Hi5 cells. Growth of the genome-modified single cells required live Hi5 feeder cells—conditioned media did not suffice— presumably because the feeder cells provide short-lived growth factors or other trophic molecules (Figure 7–figure supplement 1B). Single mCherry-positive clones developed one month after seeding and could be further grown without feeder cells. Figure 8A outlines our general strategy for establishing single-cell clones of Hi5 cells expressing fluorescent proteins.

**Figure 8.**
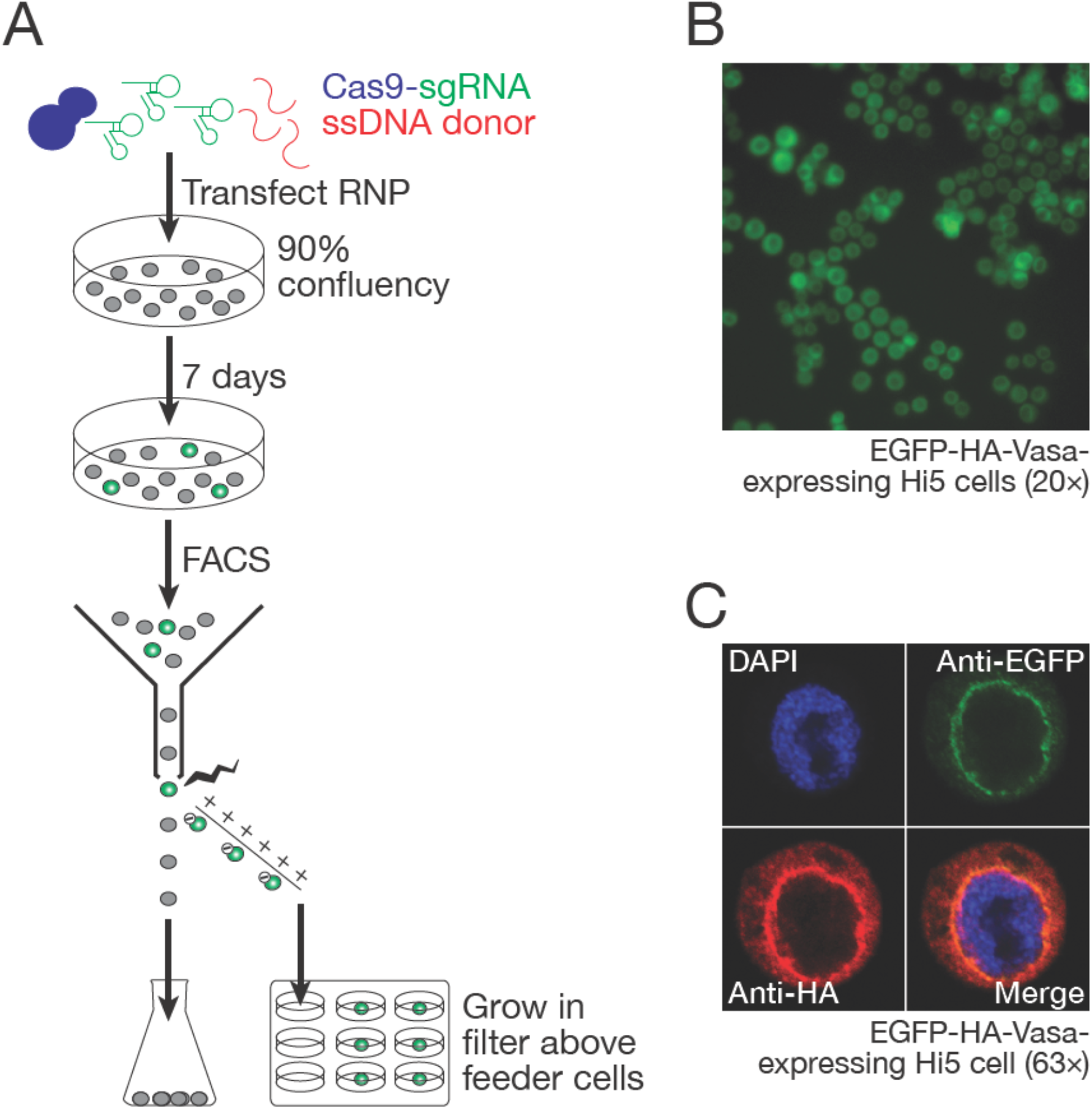
Hi5 cells contain nuage. (A) Schematic of single-clone selection of genome-edited Hi5 cells using the strategy described in Figure 7C. (B) A representative field of Hi5 cells edited to express EGFP-HA-Vasa from the endogenous locus. (C) A representative image of a fixed, EGFP-HA-Vasa-expressing Hi5 cell stained with DAPI, anti-EGFP and anti-HA antibodies. EGFP and HA staining colocalize in a perinuclear structure consistent with Vasa localizing to nuage.

To test whether an exogenous donor DNA with homologous arms could facilitate the site-specific incorporation of protein tag sequences into Hi5 genome, we designed two sgRNAs with target sites ~90 bp apart, flanking the *vasa* start codon (Figure 7C). As a donor, we used a single-stranded DNA (ssDNA) encoding EGFP and an HA epitope tag flanked by genomic sequences 787 bp upstream and 768 bp downstream of the *vasa* start codon (Figure 7C). Cas9 and the two sgRNAs were cotransfected with the ssDNA donor, and, one week later, EGFP-positive cells were detected by fluorescence microscopy. PCR amplification of the targeted region using genomic DNA from EGFP-expressing cells confirmed integration of EGFP and HA tag into the *vasa* gene (Figure 7D). Sanger sequencing further confirmed integration of EGFP and the HA tag in-frame with the *vasa* open reading frame (Supplemental file 9).

### Hi5 cell Vasa is present in a nuage-like, perinuclear structure

In the germline of *D. melanogaster* and other species, components of the piRNA biogenesis pathway, including Vasa, Aub, Ago3, and Tudor-domain proteins, localize to a perinuclear structure called nuage (*Findley SD et al., 2003*; *Lim AK* and *Kai T, 2007*; *Li C et al., 2009*; *Liu L et al., 2011*; *Webster A et al., 2015*). Vasa, a germline-specific nuage component, is widely used as a marker for nuage. In BmN4 cells, transiently transfected Vasa localizes to a perinuclear structure resembling nuage (*Xiol J et al., 2012*; *Patil AA et al., 2017*). To determine whether nuage-like structures are present in Hi5 cells, we examined Vasa localization in the Hi5 cells in which the endogenous *vasa* gene was engineered to fuse EGFP and an HA epitope tag to the Vasa amino-terminus. We used two different immunostaining strategies to detect the EGFP-HA-Vasa fusion protein: a mouse monoclonal anti-GFP antibody and a rabbit monoclonal anti-HA antibody. GFP and HA colocalized in a perinuclear structure, consistent with Vasa localizing to nuage in Hi5 cells (Figure 8C).

## Discussion

We have sequenced and assembled de novo the genome of the cabbage looper, *T. ni*, a common and destructive agricultural pest that feeds on many plants of economic importance. Examination of the *T. ni* genome and transcriptome reveals the expansion of detoxification-related gene families (Table 1 and Supplementary file 6), many members of which are implicated in insecticide resistance and are potential targets of pest control. The *T. ni* genome should enable study of the genetic diversity and population structure of this generalist pest, which adapts to different environmental niches worldwide. Moreover, as the sister order of Diptera, Lepidoptera like *T. ni* provide a counterpoint for the well-studied insect model *D. melanogaster*.

The use of Hi-C sequencing was an essential step in assembling the final 368.2 Mb *T. ni* genome into high-quality, chromosome-length scaffolds. The integration of long reads, short reads, and Hi-C provides a rapid and efficient paradigm for generating chromosome-level assemblies of other animal genomes. This strategy assembled the gene-poor and repeat-rich *T. ni* W chromosome, which is, to our knowledge, the first chromosome-level sequence of a lepidopteran W chromosome. Our analysis of autosomal, Z-linked, and W-linked transcripts provides insights into lepidopteran dosage compensation and sex determination. Our data show that *T. ni* compensates for Z chromosome dosage in the soma by reducing transcription of both Z homologs in males, but Z dosage is not compensated in the germline.

In addition to long RNAs, we characterized miRNAs, siRNAs, and piRNAs in *T. ni* gonads, soma, and cultured Hi5 cells. miRNAs are widely expressed in *T. ni* tissues, providing examples of germline-enriched and somatic miRNAs, as well as highly conserved, lepidopteran-specific, and novel *T. ni* miRNAs. Like flies, *T. ni* possess siRNAs that map to transposons, *cis*-NATs and hpRNAs. Unexpectedly, *T. ni* siRNAs— and likely all lepidopteran siRNAs—lack a 2′-*O*-methyl modification at their 3′ ends, unlike siRNAs in *D. melanogaster*. Consistent with siRNA production by a processive Dicer-2 enzyme, Hi5 cells produce phased siRNAs from the RNA genome of a latent alphanodavirus.

The commonalities and differences between *T. ni* and *D. melanogaster* small RNA pathways will help identify both deeply conserved and rapidly evolving components. A major motivation for sequencing the *T. ni* genome was the establishment of a tractable cell culture model for studying small RNAs, especially piRNAs. We believe that our genome assembly and gene-editing protocols will enable the use of *T. ni* Hi5 cells to advance our understanding of how piRNA precursors are defined and made into piRNAs and how piRNAs act to silence transposons in the germline. Hi5 cells express essentially all known piRNA pathway genes except for those specific to Drosophilids. Furthermore, *T. ni* Vasa localizes to a perinuclear, nuage-like structure in Hi5 cells, making them suitable for studying the assembly of the subcellular structures thought to organize piRNA biogenesis. We have defined genomic piRNA-producing loci in Hi5 cells, as well as in the soma, testis, and ovary. The most productive piRNA clusters are shared among ovary, testis, and Hi5 cells. In addition, Hi5 cells contain novel piRNA clusters not found in the moth itself, suggesting that the process of establishing new piRNA-producing loci can be recapitulated by experimental manipulation of Hi5 cells.

As in *D. melanogaster*, splicing of *T. ni* piRNA precursor transcripts is efficiently suppressed, yet *T. ni* lacks paralogs of the proteins implicated in splicing suppression in flies. The ability to study the mechanisms by which piRNA clusters form and how precursor RNAs are transcribed, exported, and marked for piRNA production in *T. ni* promises to reveal both conserved and lepidopteran-specific features of this pathway. Notably, the W chromosome not only is a major piRNA source, but also produces piRNAs from almost its entirety. Future studies are needed to determine whether this is a common feature of the W chromosome in Lepidoptera and other insects.

The establishment of procedures for genome editing and single-cell cloning of Hi5 cells, combined with the *T. ni* genome sequence, make this germ cell line a powerful tool to study RNA and protein function ex vivo. Our strategy combines transfection of pre-assembled Cas9/sgRNA complexes with single clone isolation using a selectable marker (e.g., EGFP) and feeder cells physically separated from the engineered cells. Compared with nucleic acid-based delivery of Cas9, transfection of Cas9 RNP minimizes the off-target mutations caused by prolonged Cas9 expression and eliminates the risk of integration of sgRNA or Cas9 sequences into the genome (*Lin S et al., 2014*; *Kim S et al., 2014*). Comparing to the use of plasmid donors (*Yu Z et al., 2014*; *Ge DT et al., 2016*), our use of ssDNA homology donors similarly reduces the chance of introducing plasmid sequences at unintended genomic sites. Techniques for injecting the embryos of other lepidopteran species have already been established (*Wang Y et al., 2013*; *Takasu Y et al., 2014*; *Zhang Z et al., 2015*). In principle, Cas9 RNP injected into cabbage looper embryos could be used to generate genetically modified *T. ni* strains both to explore lepidopteran biology and to implement novel strategies for safe and effective pest control.

## Materials and methods

### Genomic DNA libraries

Hi5 cells (ThermoFisher, Waltham, MA, USA) were cultured at 27°C in Express Five Serum Free Medium (ThermoFisher) following the manufacturer’s protocol. Thorax were dissected from four-day-old female or male T. ni pupa (Benzon Research, Carlisle, PA, USA). Cells or tissues were lysed in 2× PK buffer (200 mM Tris-HCl [pH7.5], 300 mM NaCl, 25 mM EDTA, 2% w/v SDS) containing 200 μg/ml proteinase K at 65°C for 1 h, extracted with phenol:chloroform:isoamyl alcohol (25:24:1; Sigma, St. Louis, MO, USA), and genomic DNA collected by ethanol precipitation. The precipitate was dissolved in 10 mM Tris-HCl (pH 8.0), 0.1mM EDTA, treated with 20 μg/ml RNase A at 37°C for 30 min, extracted with phenol:chloroform:isoamyl alcohol (25:24:1), and collected by ethanol precipitation. DNA concentration was determined (Qubit dsDNA HS Assay, ThermoFisher). Genomic DNA libraries were prepared from 1 µg genomic DNA (Illumina TruSeq LT kit, NextSeq 500, Illumina, San Diego, CA, USA).

Long-read genome sequencing with a 23 kb average insert range was constructed from 16 µg genomic DNA using the SMRTbell Template Prep Kit 1.0 SPv3 (Pacific Biosciences, Menlo Park, CA, USA) according to manufacturer’s protocol. Sequence analysis was performed using P6/C4 chemistry, 240 min data collection per SMRTcell on an RS II instrument (Pacific Biosciences). Mate pair libraries with 2 kb and 8 kb insert sizes were constructed (Nextera Mate Pair Library Prep Kit, Illumina) according to manufacturer’s protocol from 1 µg Hi5 cell genomic DNA. Libraries were sequenced to obtain 79 nt paired-end reads (NextSeq500, Illumina).

### Hi-C

Hi-C libraries were generated from Hi5 cells as described (Belton J-M et al., 2012), except that 50 million cells were used. Hi-C Libraries were sequenced using the NextSeq500 platform (Illumina) to obtain 79 nt, paired-end reads.

### Karyotyping

Hi5 cells were first incubated in Express Five medium containing 1 µg/ml colcemid at 27°C for 8 h (Schneider I, 1979), then in 4 ml 0.075 M KCl for 30 min at 37°C, and fixed with freshly prepared methanol:acetic acid (3:1, v:v) precooled to -20°C. Mitotic chromosomes were spread, mounted by incubation in ProLong Gold Antifade Mountant with DAPI (4ʹ,6ʹ-diamidino-2-phenylindole) (ThermoFisher) overnight in the dark, and imaged using a DMi8 fluorescence microscope equipped with an 63× 1.40 N.A. oil immersion objective (HCX PL APO CS2, Leica Microsystems, Buffalo Grove, IL, USA) as described (Matijasevic Z et al., 2008).

### Small RNA libraries

Ovaries, testes, and thoraces were dissected from cabbage looper adults 24–48 h after emerging. Total RNA (30 µg) was isolated (mirVana miRNA isolation kit, Ambion, Austin, TX, USA) and sequenced using the NextSeq500 platform (Illumina) to obtain 59 nt single-end reads as previously described (Han BW et al., 2015b).

### RNA-seq

Adult ovaries, testes, or thoraces were dissected from cabbage looper adults 24 to 48 h after emerging. Total RNA (3 µg) was purified (mirVana miRNA isolation kit, Ambion) and sequenced as described (Zhang Z et al., 2012) using the NextSeq500 platform (Illumina) to obtain 79 nt, paired-end reads.

### Genome assembly

Canu v1.3 (Koren S et al., 2017) was used to assemble long reads into contigs, followed by Quiver (github.com/PacificBiosciences/GenomicConsensus) to polish the contigs using the same set of reads. Pilon (Walker BJ et al., 2014) was used to further polish the assembly using Illumina paired-end reads. Finally, to assemble the genome into chromosome-length scaffolds, we joined the contigs using Hi-C reads and LACHESIS (Burton JN et al., 2013). The mitochondrial genome was assembled separately using MITObim (six iterations, D. melanogaster mitochondrial genome as bait; [Hahn C et al., 2013]).

To evaluate the quality of the genome assembly, we ran BUSCO v3 (Simão FA et al., 2015) using the arthropod profile and default parameters to identify universal single-copy orthologs. We further evaluated genome quality using conserved gene sets: OXPHOS and CRP genes. *B. mori* and *D. melanogaster* OXPHOS and CRP protein sequences were retrieved (Marygold SJ et al., 2007; Porcelli D et al., 2007) and BLASTp was used to search for their T. ni homologs, which were further validated by querying using InterPro (Jones P et al., 2014; Mitchell A et al., 2015). We also assembled T. ni genomes from male and female animals respectively using SOAPdenovo2 (kmer size 69; [Luo R et al., 2012]). We then compared the animal genomes with the T. ni genome assembled from Hi5 cells using QUAST (-m 500; [Gurevich A et al., 2013] and the nucmer and mummerplot (–layout–filter) functions from MUMmer 3.23 (Kurtz S et al., 2004).

### Genome annotation

To annotate the T. ni genome, we first masked repetitive sequences and then integrated multiple sources of evidence to predict gene models. We used RepeatModeler to define repeat consensus sequences and RepeatMasker (-s -e ncbi) to mask repetitive regions (Smit AFA et al., 2017). We used RNAmmer (Lagesen K et al., 2007) to predict 8S, 18S, 28S rRNA genes, and Barrnap (https://github.com/tseemann/barrnap) to predict 5.8S rRNA genes. We used Augustus v3.2.2 (Stanke M et al., 2006) and SNAP (Korf I, 2004) to computationally predicted gene models. Predicted gene models were compiled by running six iterations of MAKER (Campbell MS et al., 2014), aided with homology evidence of well annotated genes (UniProtKB/Swiss-Prot and Ensembl) and of transcripts from related species (*B. mori* [Suetsugu Y et al., 2013] and *D. melanogaster* [Attrill H et al., 2016]). We used BLAST2GO (Conesa A et al., 2005) to integrate results from BLAST, and InterPro (Mitchell A et al., 2015) to assign GO terms to each gene. We used MITOS (Bernt M et al., 2013) web server to predict mitochondrial genes and WebApollo (Lee E et al., 2013) for manual curation of genes of interest. To characterize telomeres, we used (TTAGG)200 (Robertson *HM* and *Gordon* KHJ, 2006) as the query to search the T. ni genome using BLASTn with the option ‘-dust no’ and kept hits longer than 100 nt. The genomic coordinates of these hits were extended by 10 kb to obtain the subtelomeric region.

### Orthology and evolution

To place genes into ortholog groups, we compared the predicted proteomes from 21 species (Supplementary file 5). Orthology assignment was determined using OrthoMCL (Hirose *Y and Manley* JL, 1997) with default parameters. MUSCLE v3.8.31 (Edgar RC, 2004) was used for strict 1:1:1 orthologs (n = 381) to produce sequence alignments. Conserved blocks (66,044 amino acids in total) of these alignments were extracted using Gblocks v0.91b (Castresana J, 2000) with default parameters, and fed into PhyML 3.0 (Vastenhouw NL et al., 2010) (maximum likelihood, bootstrap value set to 1000) to calculate a phylogenetic tree. The human and mouse predicted proteomes were used as an outgroup to root the tree. The tree was viewed using FigTree (http://tree.bio.ed.ac.uk/software/figtree/) and iTOL (Shirayama M et al., 2012).

### Sex determination and sex chromosomes

To identify sex-linked contigs, we mapped genomic sequence reads from males and females to the contigs. Reads with MAPQ scores ≥20 were used to calculate contig coverage, which was then normalized by the median coverage. The distribution of normalized contig coverage ratios (male:female ratios, M:F ratios) was manually checked to empirically determine the thresholds for Z-linked and W-linked contigs (M:F ratio >1.5 for Z-linked contigs and M:F ratio < 0.5 for W-linked contigs). Lepidopteran masc genes were obtained from Lepbase (Challis RJ et al., 2016). Z/AA ratio was calculated according to (Gu L et al., 2017).

### Gene families for detoxification and chemoreception

To curate genes related to detoxification and chemoreception, we obtained seed alignments from Pfam (Finn RD et al., 2016) and ran hmmbuild to build HMM profiles of cytochrome P450 (P450), amino-and carboxy-termini of glutathione-S-transferase (GST), carboxylesterase (COE), ATP-binding cassette transporter (ABCs), olfactory receptor (OR), gustatory receptor (GR), ionotropic receptor (IR), and odorant binding (OBP) proteins, (Supplementary file 6, 7 and 8). We then used these HMM profiles to search for gene models in the predicted T. ni proteome (hmmsearch, e-value cutoff: 1 × 10 - 5). We also retrieved reference sequences of P450, GST, COE, ABC, OR, GR, IR, OBP, and juvenile hormone pathway genes from the literature (Hekmat-Scafe DS et al., 2002; ; Xavier B et al., 2005; Wanner KW and Robertson HM, 2008; Yu Q et al., 2008; Benton R et al., 2009; Gong D-P et al., 2009; Yu Q-Y et al., 2009; Croset V et al., 2010; Ai J et al., 2011; Liu S et al., 2011; Dermauw W and Van Leeuwen T, 2014; van Schooten B et al., 2016). These were aligned to the T. ni genome using tBLASTx (Altschul SF et al., 1990) and Exonerate (Slater GSC and Birney E, 2005) to search for homologs. Hits were manually inspected to ensure compatibility with RNA-seq data, predicted gene models, known protein domains (using CDD [Marchler-Bauer A et al., 2015]) and homologs from other species. P450 genes were submitted to David Nelson’s Cytochrome P450 Homepage (Nelson DR, 2009) for nomenclature and classification. Sequences and statistics of these genes are in Supplementary files 6, 7 and 8.

To determine the phylogeny of these gene families, we aligned the putative protein sequences from *T. ni* and *B. mori* genomes using MUSCLE (Edgar RC, 2004), trimmed the multiple sequence alignments using TrimAl (Capella-Gutiérrez S et al., 2009) (with the option -automated1), and performed phylogenetic analysis (PhyML 3.0 [Vastenhouw NL et al., 2010], with parameters: -q –datatype aa –run_id 0 – no_memory_check -b -2). Phylogenetic trees were visualized using FigTree (http://tree.bio.ed.ac.uk/software/figtree/).

To curate opsin genes, we used opsin mRNA and peptide sequences from other species (Zimyanin VL et al., 2008; Futahashi R et al., 2015) to search for homologs in T. ni. To discriminate opsin genes from other G-protein-coupled receptors, we required that the top hit in the NCBI nr database and Uniprot were opsins.

### Transposon analysis

To determine transposon age, we calculated the average percent divergence for each transposon family: the percent divergence (RepeatMasker) of each transposon copy was multiplied by its length, and the sum of all copies were divided by the sum of lengths of all copies in the family (Pace JK and Feschotte C, 2007). We used TEMP (Zhuang J et al., 2014) to identify transposon insertions in the Hi5 genome.

### miRNA and siRNA analysis

mirDeep2 (Friedländer MR et al., 2008; Friedlander MR et al., 2012) with default parameters predicted miRNA genes. Predicted miRNA hairpins were required to have homology (exact seed matches and blastn e-value < 1 × 10-5) to known miRNAs and/or miRDeep2 scores ≥10. miRNAs were named according to exact seed matches and high sequence identities (blastn e-value < 1 × 10-5) with known miRNA hairpins.

To determine the conservation status of T. ni miRNAs, putative T. ni miRNAs were compared with annotated miRNAs from *A. aegypti, A. mellifera, B. mori, D. melanogaster, H. sapiens, M. musculus, M. sexta, P. xylostella, and T. castaneum*: conserved miRNAs were required to have homologous miRNAs beyond Lepidoptera. To compare siRNA abundance in oxidized and unoxidized small RNA-seq libraries, we normalized siRNA read counts to piRNA cluster-mapping reads (piRNA cluster read counts had >0.98 Pearson correlation coefficients between oxidized and unoxidized libraries in all cases). Because piRNA degradation products can be 20–22 nt long, we excluded potential siRNA species that were prefixes of piRNAs (23–35 nt). To search for viral transcripts in T. ni, we downloaded viral protein sequences from NCBI (http://www.ncbi.nlm.nih.gov/genome/viruses/) and used using tBLASTn to map them to the T. ni genome and to the transcriptomes of Hi5 cells and five T. ni tissues. We filtered hits (percent identity ≥ 0.80, e-val ≤ 1 × 10-20, and alignment length ≥ 100) and mapped small RNA-seq reads to the identified viral transcripts. Candidate genomic hairpins were defined according to (Okamura K et al., 2008b). Candidate cis-NATs were defined according to (Ghildiyal M et al., 2008).

### piRNA analysis

To determine the genomic coordinates of piRNA-producing loci, we mapped small RNAs to the genome as described (Han BW et al., 2014). We then calculated the abundance of piRNAs in 5 kb genomic windows. For each window, we counted the number of uniquely mapped reads and the number of reads mapped to multiple loci (multimappers) by assigning reads using an expectation-maximization algorithm. Briefly, each window had the same initial weight. The weight was used to linearly apportion multimappers. During the expectation (E) step, uniquely mapped reads were unambiguously assigned to genomic windows; multimappers were apportioned to the genomic windows they mapped to, according to the weights of these windows. At the maximization (M) step, window weights were updated to reflect the number of reads each window contained from the E step. The E and M steps were run iteratively until the Manhattan distance between two consecutive iterations was smaller than 0.1% of the total number of reads.

To identify differentially expressed piRNA loci, we used the ppm and rpkm values, normalized to the total number of uniquely mapped reads, to measure piRNA abundance. For analyses including all mapped reads (uniquely mapped reads and multimappers), reads were apportioned by the number of times that they were mapped to the genome. To make piRNA loci comparable across tissues, we merged piRNA loci from ovary, testis, female and male thorax, and Hi5 cells. For the comparison between female and male thoraces, the cluster on tig00001980 was removed as this cluster likely corresponds to a mis-assembly. We used Spearman correlations to calculate the pairwise correlations of piRNA abundances. As for defining sex-linked contigs, we calculated M:F ratios and used the same thresholds to determine whether a piRNA cluster was sex-linked. A piRNA locus was considered to be differentially expressed if the ratio between the two tissues was >2 or <0.5 and FDR <0.1 (after t-test). Splice sites were deemed to be supported by RNA-seq data when supported by at least one data set. We used AUGUSTUS (Stanke M et al., 2006), with the model trained for T. ni genome-wide gene prediction, to predict gene models and their splice sites in T. ni piRNA clusters.

### β-elimination

Total RNAs were extracted from Hi5 cells using mirVana kit as described previously. We then incubated 100 µg total RNA with 25 mM NaIO4 in borate buffer (148 mM Borax,148 mM Boric acid, pH 8.6) for 30 min at room temperature, beta-elimination was performed in 50 mM NaOH at 45°C for 90 min (Horwich MD et al., 2007). The resultant RNA was collected by ethanol precipitation.

### sgRNA design

sgRNAs for the target loci (5′-end of ciwi and 5′-end of vasa) were designed using crispr.mit.edu (Hsu PD et al., 2013) to retrieve all possible guide sequences, and guide sequences adjacent to deletion or insertion targets were chosen. Supplementary file 9 lists guide sequences.

### ssDNA donor purification

Donor template sequence was produced as a gBlock (Integrated DNA Technologies, San Diego, CA, USA). A biotinylated forward primer and a standard reverse primer were used in PCR to generate a double-stranded, biotinylated DNA donor. The biotinylated DNA was captured on M-280 streptavidin Dynabeads (ThermoFisher), and the biotinylated strand was separated from the non-biotinylated strand essentially as described in the manufacturer’s protocol. Supplemental file 10 provides a detailed protocol.

### Transfection of Hi5 cells

sgRNAs were transcribed using T7 RNA polymerase, gel purified, then incubated with Cas9 in serum-free Hi5 culture medium supplemented with 18 mM l-glutamine. The resulting sgRNA/Cas9 RNPs were incubated with Trans-IT insect reagent (Mirus Bio, Madison, WI, USA) for 15 min at room temperature, then evenly distributed onto 90% confluent Hi5 cells. Culture medium was replaced with fresh medium 12 h later. Genomic DNA was isolated and analyzed by PCR 48 h later.

### PCR to validate genomic editing in transfected cells

Forty eight hours after transfection, Hi5 cells from one 90% confluent well of a six-well plate (Corning, Corning, NY, USA) were collected, washed once with PBS (ThermoFisher) and lysed in 2× PK buffer containing 200 μg/ml proteinase K, extracted with phenol:chloroform:isoamyl alcohol (25:24:1), and then genomic DNA collected by ethanol precipitation. Deletions in ciwi were detected by PCR using primers flanking the deleted region (Supplementary file 9). To confirm deletions by sequencing PCR products were resolved by agarose gel electrophoresis, purified (QIAquick Gel Extraction Kit, QIAGEN, Germantown, MD, USA), and cloned into pCR-Blunt II-Topo vector (ThermoFisher). The recombinant plasmid was transformed into Top10 competent E.coli (ThermoFisher) following supplier’s protocol. PCR products amplified using M13 (-20) forward and M13 reverse primers from a sample of a single bacterial colony were sequenced by GENEWIZ (South Plainfield, NJ, USA).

### Single clone selection

Wild-type Hi5 cells were seeded into a 96-well Transwell permeable support receiver plate (Corning, Corning, NY, USA) at 30% confluence and incubated overnight in serum free medium with 100 U/ml penicillin and 100 μg/ml streptomycin. A Transwell permeable support insert plate with media in each well was inserted into the receiver plate, and a single EGFP-positive cell was sorted into each insert well by FACS. After 14 days incubation at 27°C, wells were examined for EGFP-positive cell clones using a DMi8 fluorescent microscope (Leica).

### Immunostaining

EGFP-HA-Vasa-expressing Hi5 cells were seeded on 22 x 22 mm cover slips (Fisher Scientific, Pittsburgh, PA, USA) in a well of a six-well plate (Corning). After cells had attached to the coverslip, the medium was removed and cells were washed three times with PBS (Gibco). Cells were fixed in 4% (w/v) methanol-free formaldehyde (ThermoFisher) in PBS at room temperature for 15 min, washed three times with PBS, permeabilized with 0.1% (w/v) Triton X-100 in PBS for 15 min at room temperature, and then washed three times with PBS. For antibody labeling, cells were incubated in 0.4% (v/v) Photo-Flo in 1× PBS for 10 min at room temperature, then 10 min in 0.1% (w/v) Triton X-100 in PBS and 10 min in 1× ADB-PBS (3g/l bovine serum albumen, 1% (v/v) donkey serum, 0.005% (w/v) Triton X-100 in 1× PBS). Next, cells were incubated with primary antibodies (mouse anti-GFP antibody (GFP-1D2, Developmental Studies Hybridoma Bank, Iowa City, IA, USA) and rabbit anti-HA Tag antibody (C29F4, Cell Signaling, Danvers, MA, USA), diluted 1:200 in ADB (30g/l BSA, 10% (v/v) donkey serum, 0.05% (w/v) Triton X-100 in 1× PBS) at 4°C overnight. After three washes in PBS, cells were incubated sequentially in 0.4% (v/v) Photo-Flo in 1× PBS, 0.1% (w/v) Triton X-100 in PBS, and 1× ADB-PBS, each for 10 min at room temperature. Cells were then incubated with secondary Alexa Fluor 488-labeled donkey anti-mouse (ThermoFisher) and Alexa Fluor 680-labeled donkey anti-rabbit (ThermoFisher) antibodies, diluted 1:500 in ADB at room temperature for one hour. After washing three times with 0.4% (v/v) Photo-Flo in 1× PBS and once with 0.4% (v/v) Photo-Flo in water, coverslips were air dried in the dark at room temperature. Slides were mounted in ProLong Gold Antifade Mountant with DAPI and examined by confocal microscopy (TCS SP5 II Laser Scanning Confocal, Leica).

### Data deposition

The T. ni Whole Genome Shotgun project has been deposited at DDBJ/ENA/GenBank under the accession NKQN00000000. The version described here is version NKQN01000000. All sequencing data are available through the NCBI Sequence Read Archive under the accession number PRJNA336361. Further details are available at the Cabbage Looper Database (http://cabbagelooper.org/).

## Acknowledgements

We thank members of the Weng and Zamore laboratories for helpful discussions and comments on the manuscript; UMass Deep Sequencing Core for Pacific Biosciences sequencing; Zdenka Matijasevic for sharing the karyotyping protocol. This work was supported in part by National Institutes of Health grants R37GM062862 to PDZ and HD078253 to ZW.

## Author contributions

YF, Assembly strategy, Bioinformatics analysis, Writing; YY, Design and preparation of RNA-seq, small RNA-seq, Hi-C libraries, Writing; HZ, Development of Hi5 genome editing and cloning procedures; GF, Cabbage looper breeding; JW, DNA-seq; KAQ, *P*. *xylostella* small RNA-seq; ZW, Project design and management, Assembly strategy, Writing; PDZ, Project conception, design and management, Assembly strategy, Writing.

---

**Figure 1—figure supplement 1.**
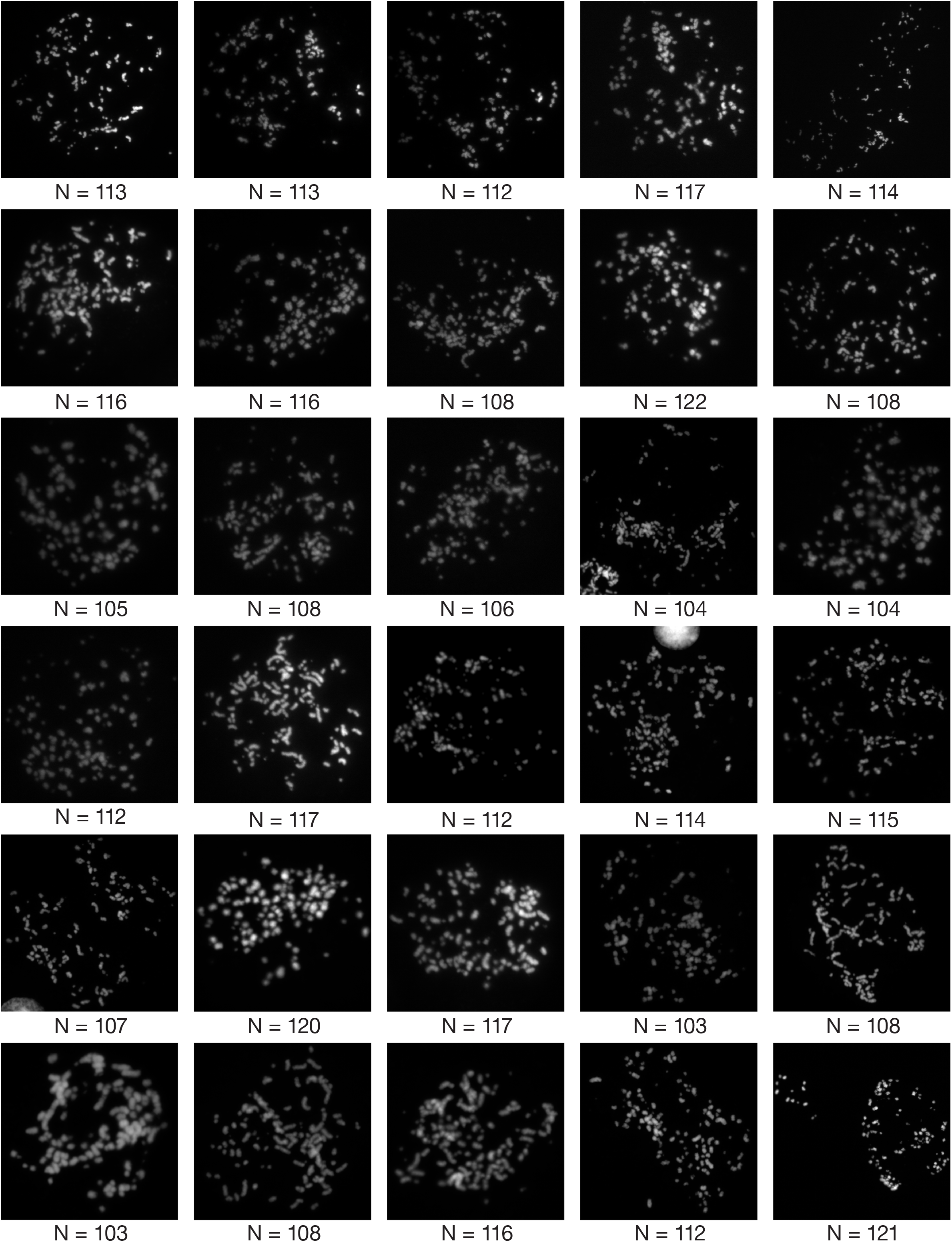
Hi5 cell Karyotyping. Thirty images showing the numbers of chromosomes (N) in Hi5 cells. N ranged from 103 to 122; mean ± S.D. = 111.7 ± 5.45. Since lepidopteran cell lines are typically tetraploid, the haploid genome likely contains 28 (mean ± S.D. = 27.9 ± 1.36) pairs of chromosomes.

**Figure 1—figure supplement 2.**
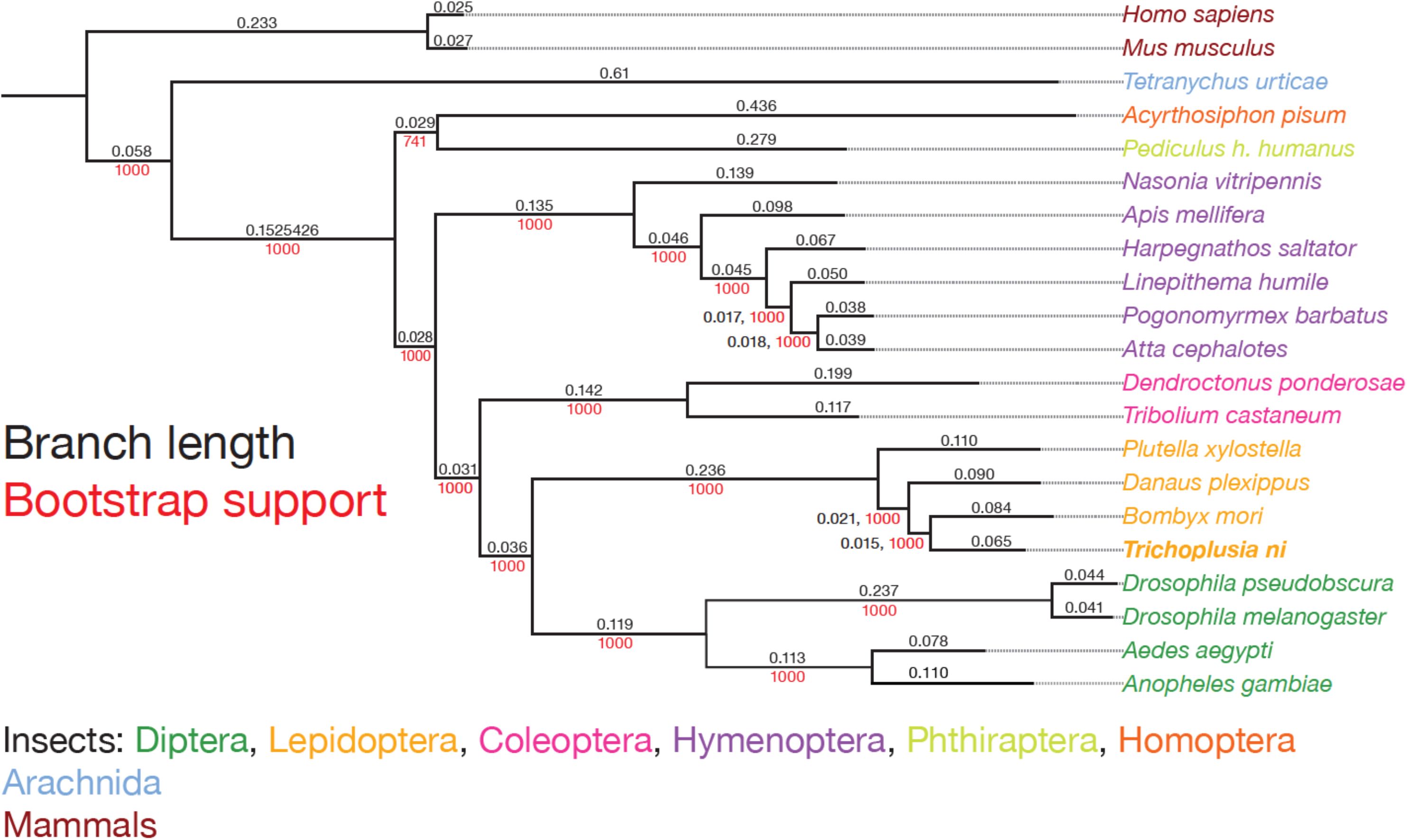
Phylogenetic tree of 21 species showing the scale, branch lengths and bootstrap support. Strict 1:1:1 orthologs were used to compute the phylogenetic tree using the maximum likelihood method. Black, branch length; red, bootstrap support.

**Figure 1—figure supplement 3.**
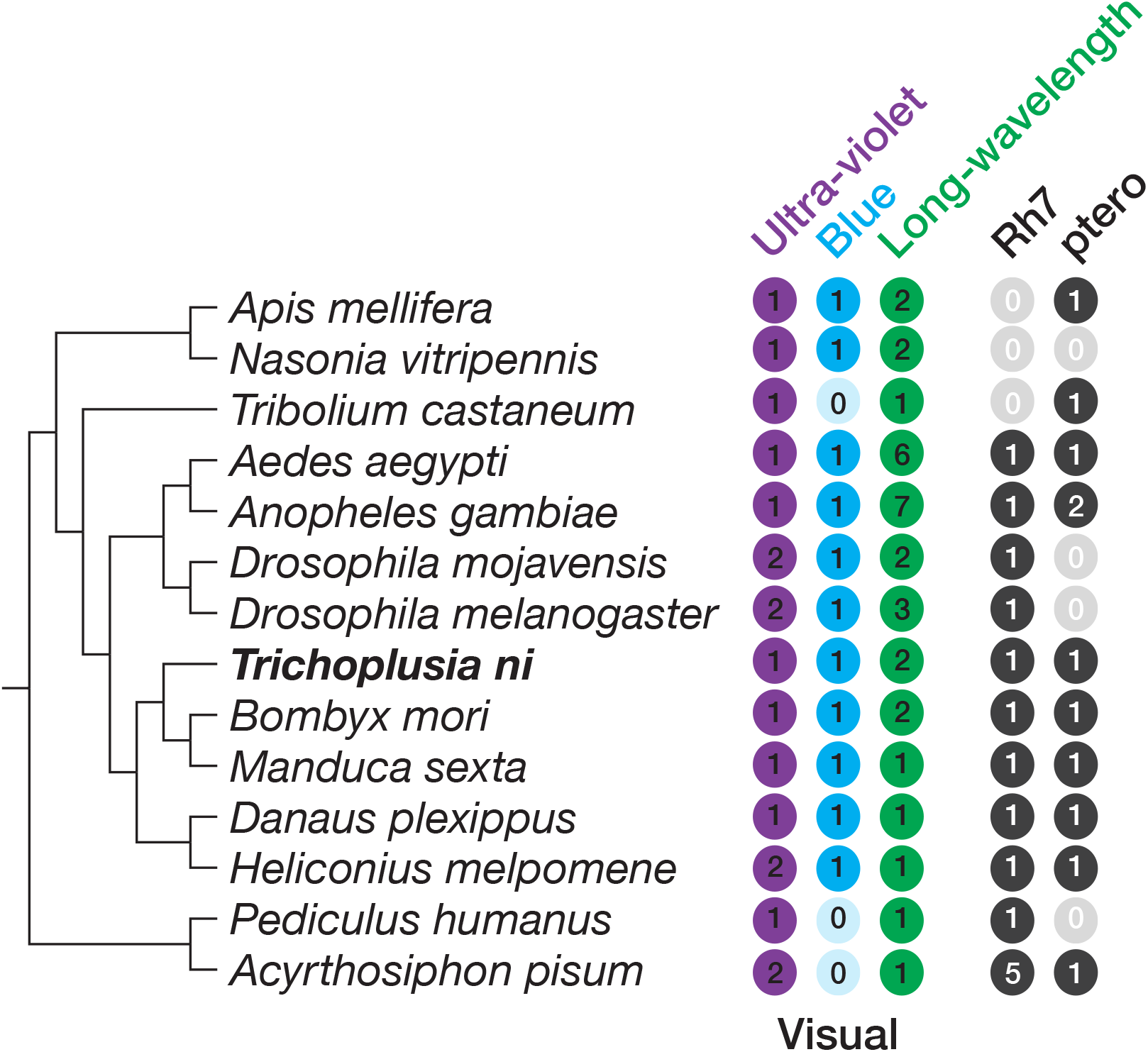
Opsins in insects.

**Figure 2—figure supplement 1.**
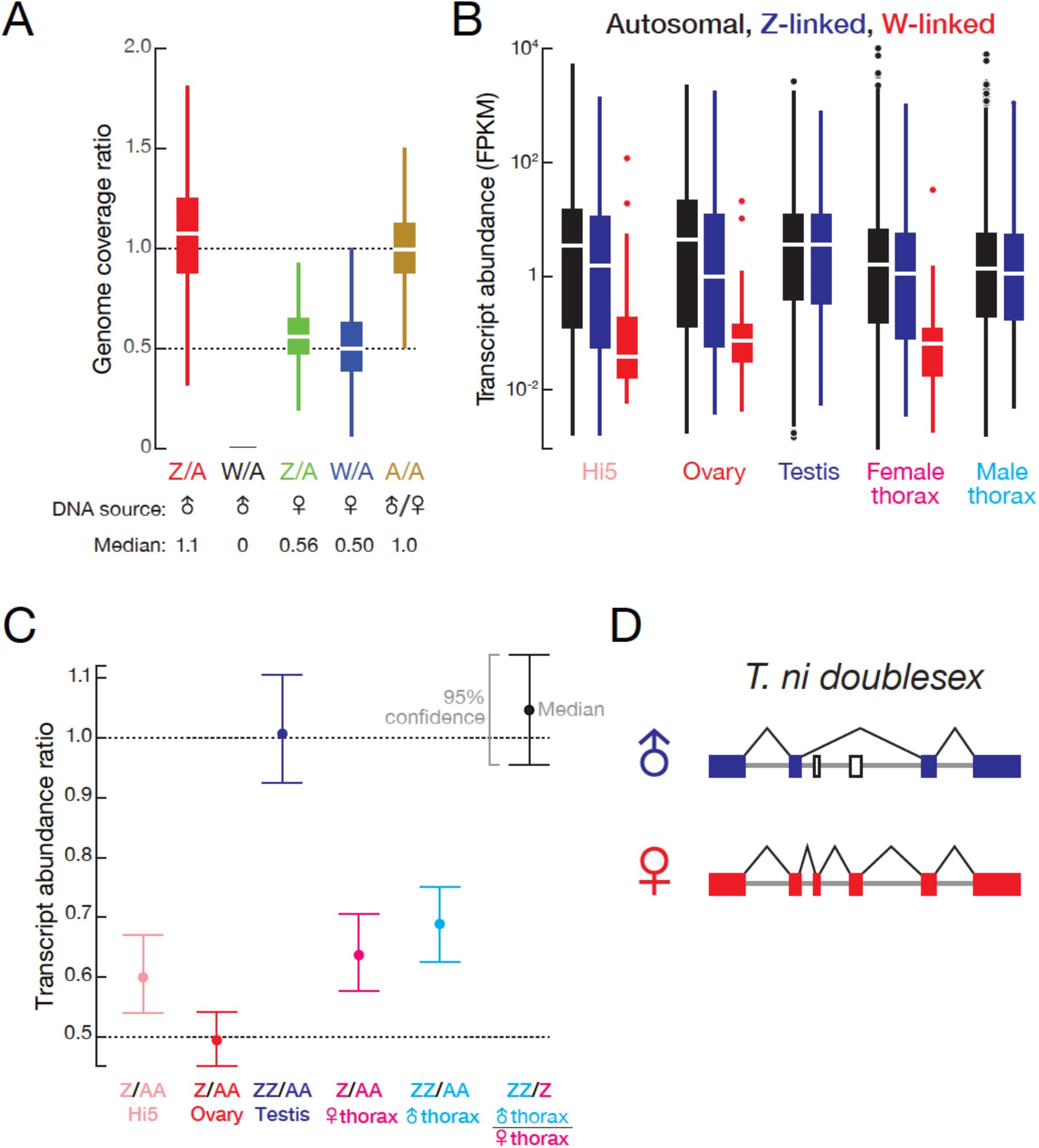
(A) Genomic coverage comparison of Z-linked, W-linked and autosomal contigs. Contig coverage was shuffled 1,000,000 times to calculate the coverage ratio. Outliers are not shown. (B) Autosomal, Z-linked and W-linked transcript abundance in Hi5 cells and *T. ni* tissues. (C) Transcript abundance ratios of autosomal, Z-linked, and W-linked genes in Hi5 cells and *T. ni* tissues. Error bars represent 95% confidence interval estimated from 1,000 bootstrap replicates. (D). Sex-specific splicing of *T. ni doublesex* pre-mRNA.

**Figure 2—figure supplement 2.**
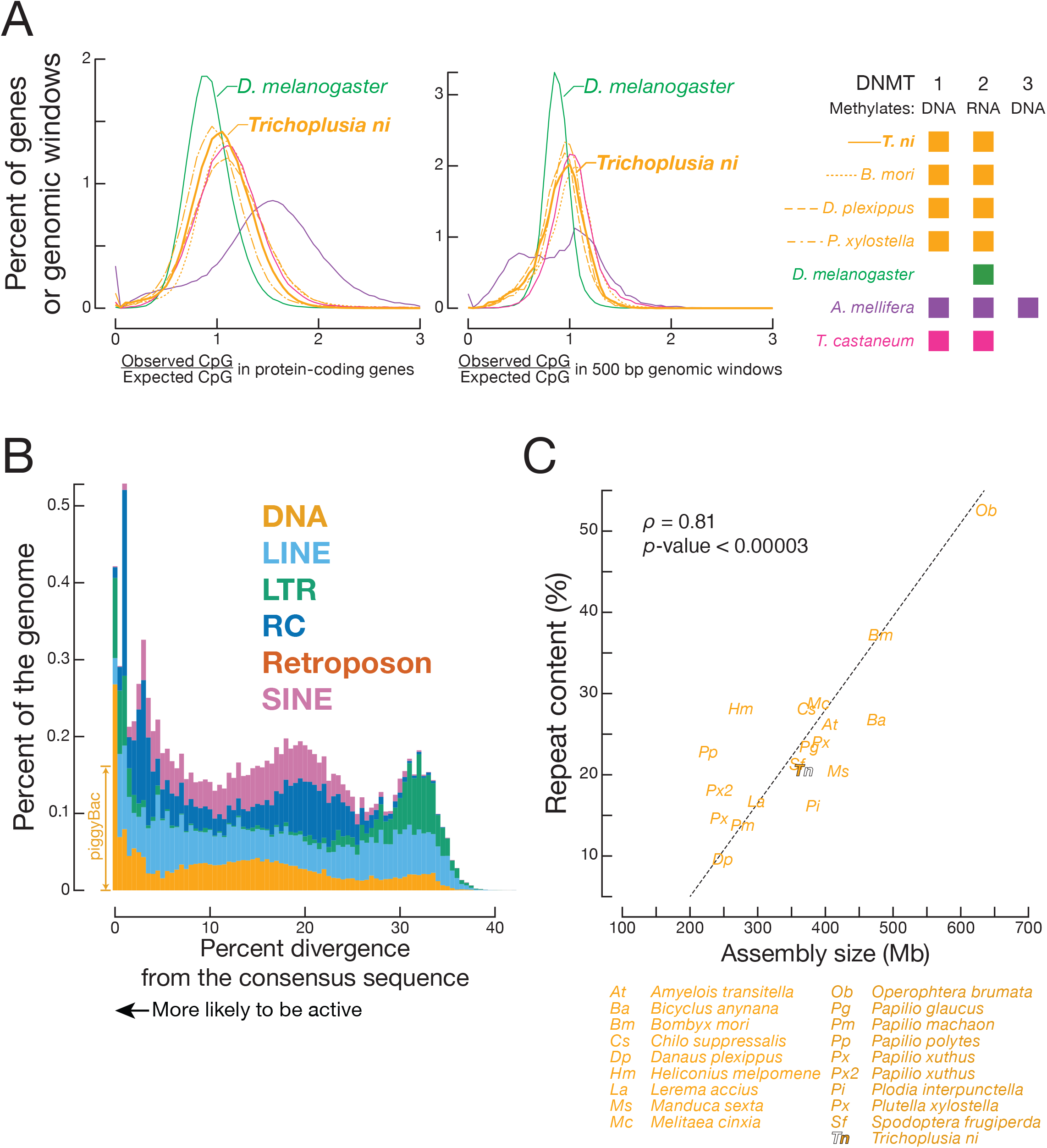
CpG ratios and transposons. (A) Distribution of observed-to-expected CpG ratios in protein-coding genes (left panel) and in 500 bp genomic windows (right panel) in *A. mellifera, B. mori, D. plexippus, D. melanogaster, P. xylostella, T. castaneum*, and *T. ni*. (B) Proportion of the genome occupied by transposons versus transposon sequence divergence. Sequence divergence was calculated by comparing individual transposon copies with the corresponding consensus sequence (Materials and methods). (C) Repeat content in lepidopteran genomes.

**Figure 4—figure supplement 1.**
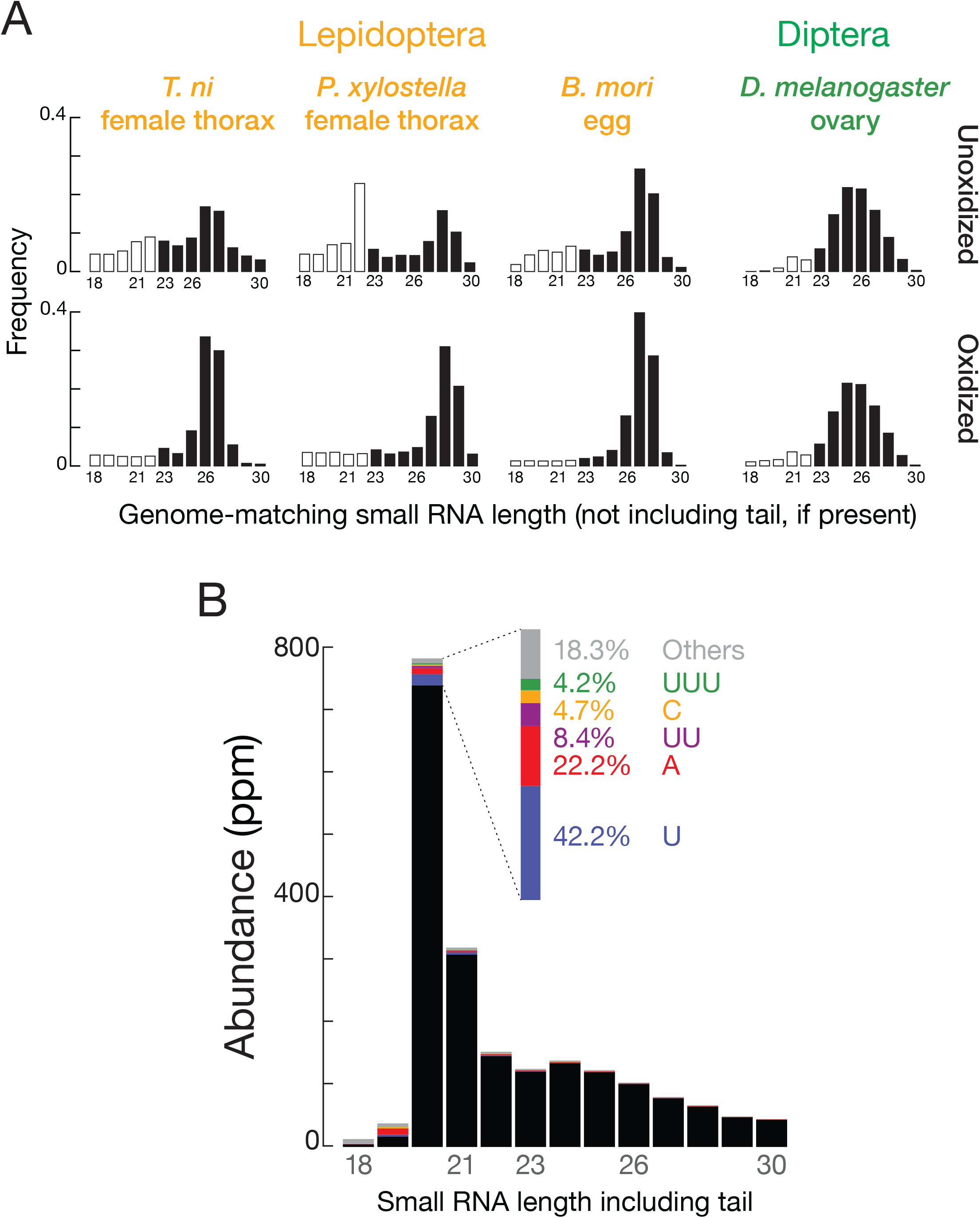
(A) siRNA length distributions for multiple insects in oxidized and unoxidized small RNA-seq libraries. (B) Length distribution of fully matched and tailed TNCL virus-siRNAs.

**Figure 5—figure supplement 1.**
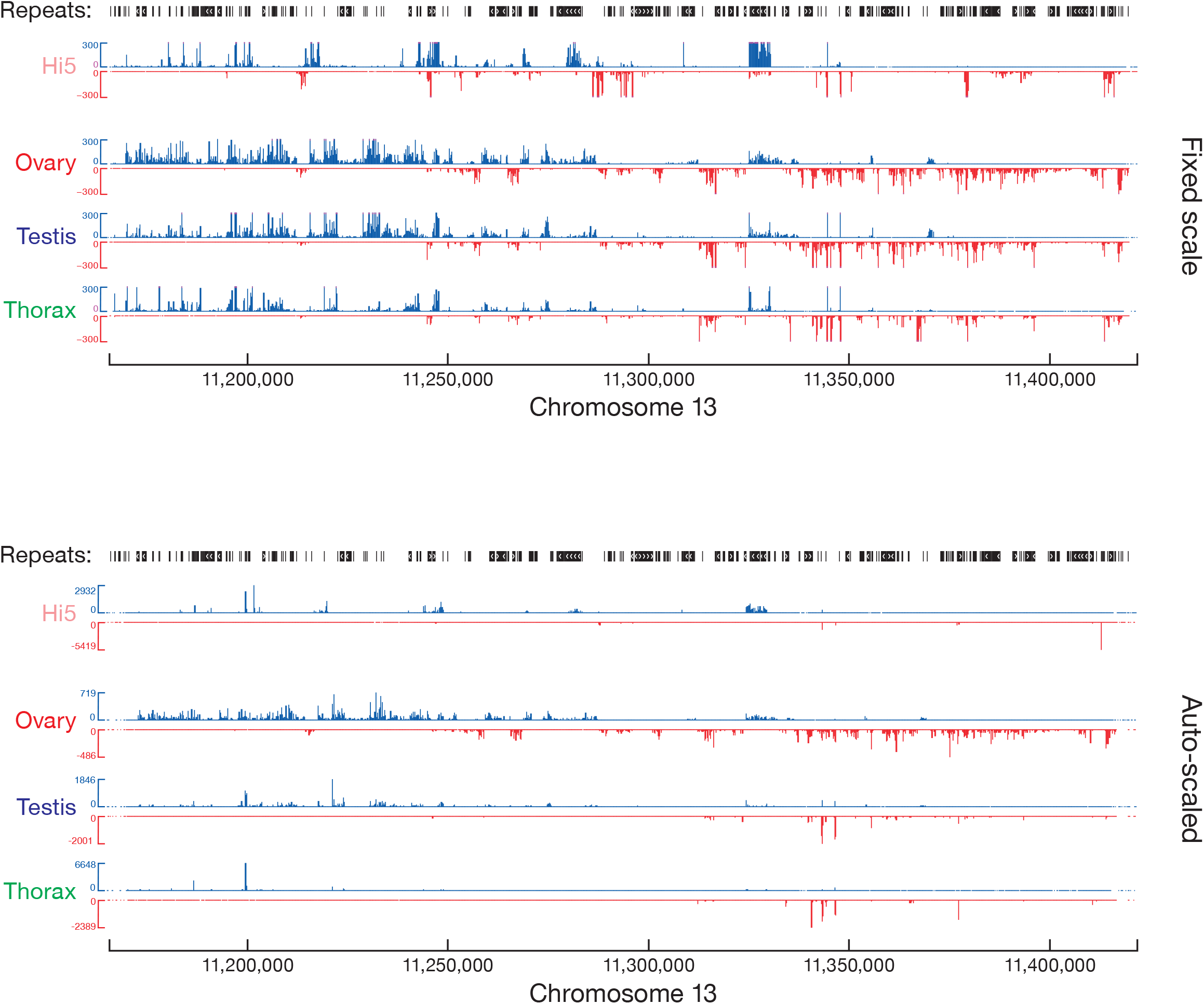
piRNA abundance (ppm) along the most productive piRNA cluster. Top, fixed scale (some data clipped); bottom, auto-scaled.

**Figure 5—figure supplement 2.**
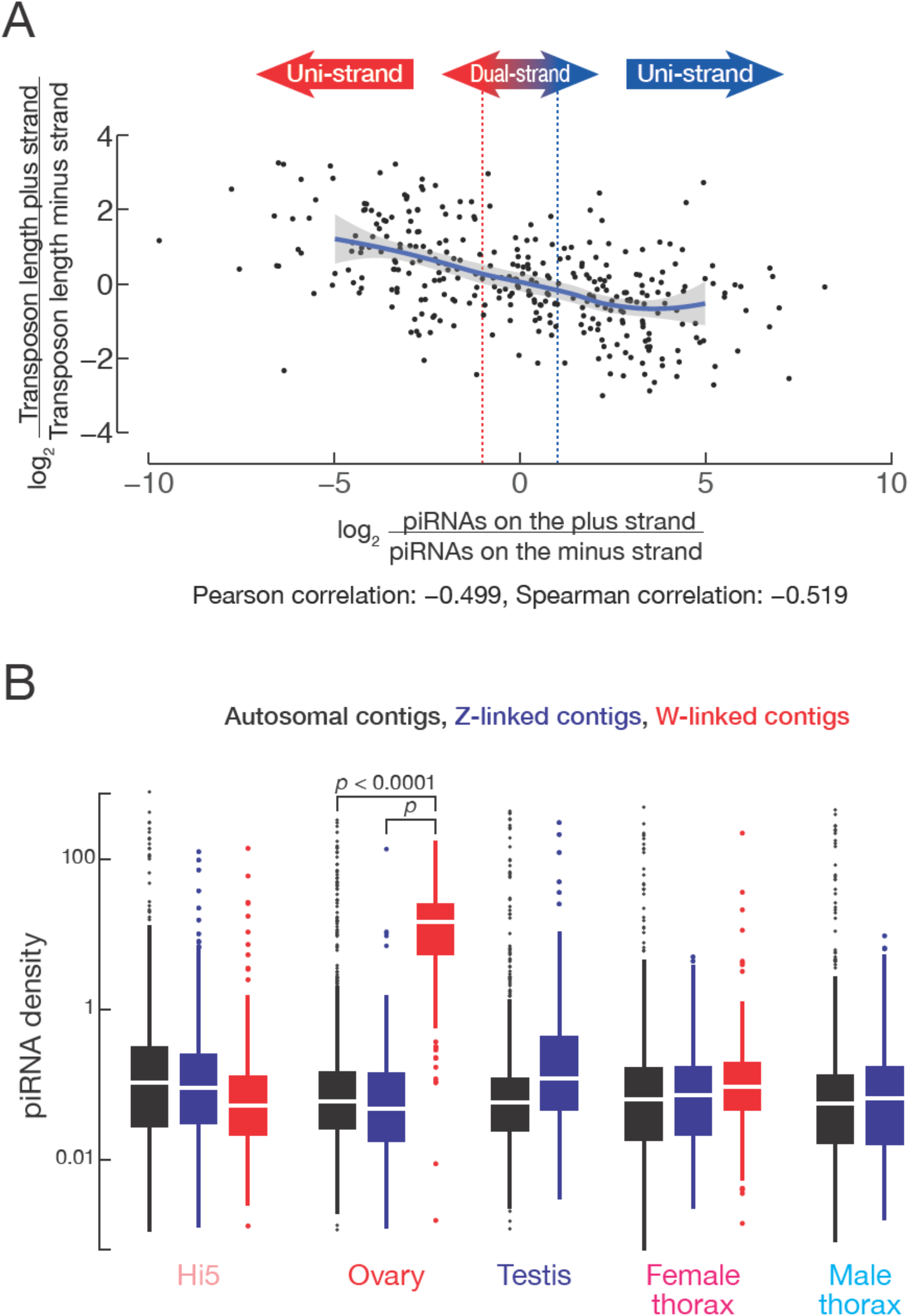
(A) piRNA clusters tend to produce piRNAs that are antisense to transposons. The *x*-axis represents the ratio of piRNAs from the plus strand over piRNAs from the minus strand, with the dotted lines indicating twofold difference. The *y*-axis indicates the ratio of transposons lengths on the plus strand over transposon length on the minus strand. The solid line indicates regression line and shading indicates 95% confidence interval by LOWESS. (B) piRNA densities on autosomal, Z-linked and W-linked contigs in Hi5 cells, ovary, testis, and female and male thorax.

**Figure 6—figure supplement 1.**
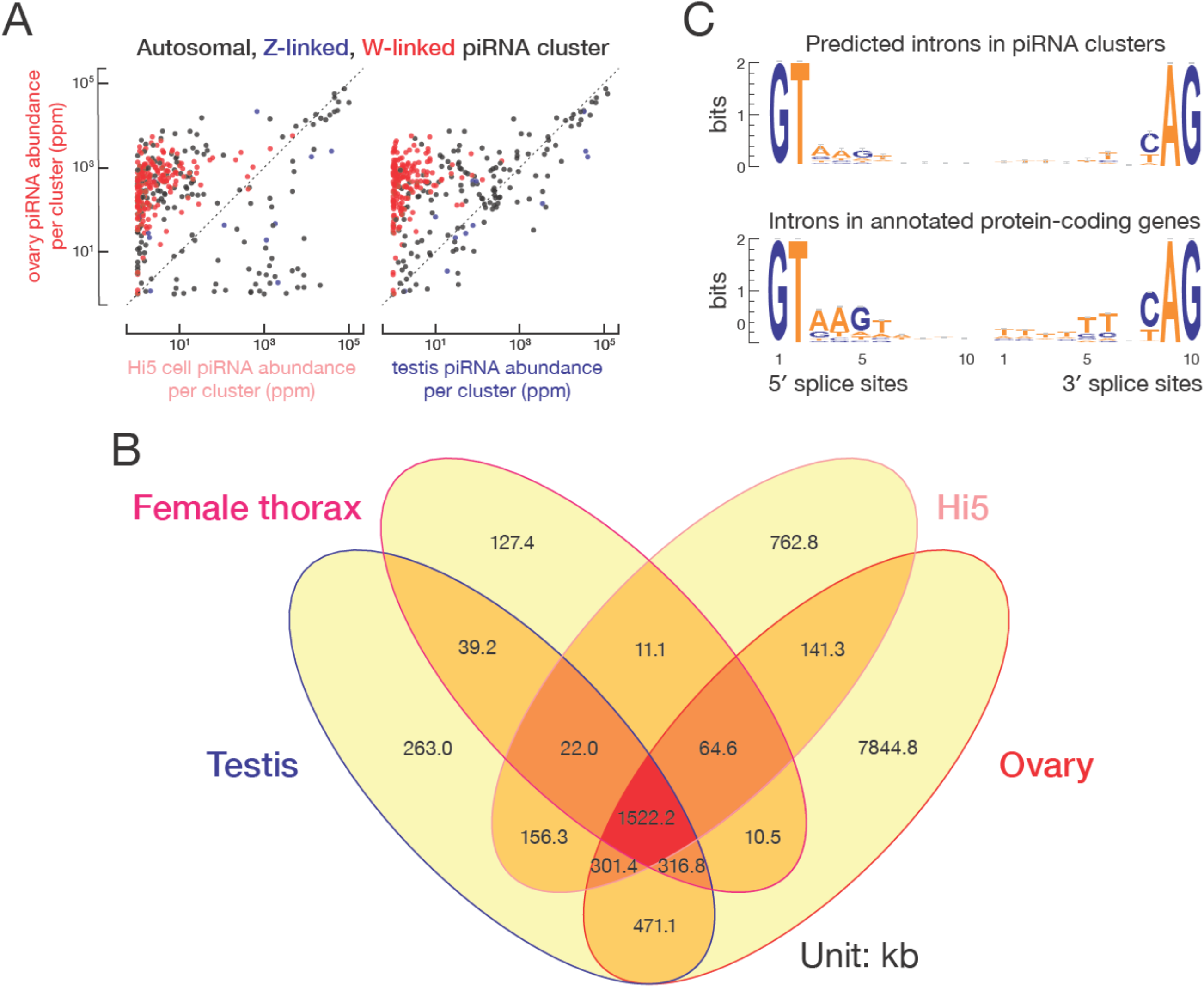
(A) Comparison of piRNA abundance (ppm) from ovary and Hi5 piRNA-producing loci and of piRNA abundance (ppm) from ovary and testis piRNA-producing loci. (B) piRNA cluster lengths in *T. ni* ovary, testis, thorax, and Hi5 cells. (C) Motifs around intron boundaries of predicted protein-coding gene models within and outside of piRNA clusters.

**Figure 7—figure supplement 1.**
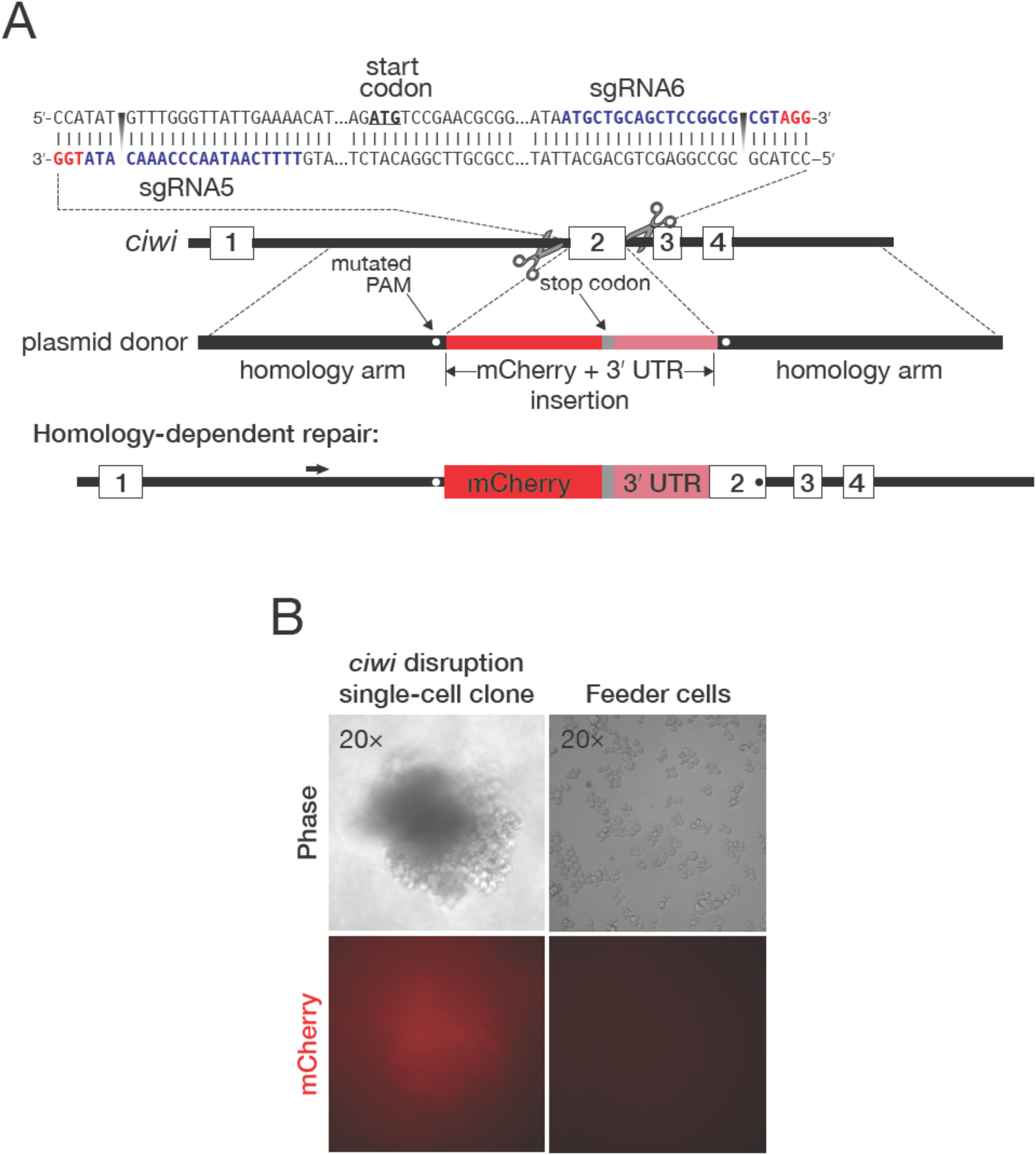
(A) Strategy for disrupting *ciwi* by inserting sequence encoding mCherry 5′ to the *ciwi* start codon. Red, protospacer-adjacent motif (PAM); blue, protospacer sequence. Arrows indicate the diagnostic forward and reverse primers used in PCR to detect genomic deletions. (B) A representative mCherry-disrupted *ciwi* clone derived from a single cell and growing as a ball of cells in a Transwell permeable support insert. Feeder Hi5 cells grown below the permeable support in the receiver well are shown for comparison.

## Supplementary Files

**Supplementary file 1**. *T. ni* genome statistics. (A) BUSCO assessments of *T. ni* and six other genomes. (B) CRP genes. (C) Genes in the OXPHOS pathway. (D) Genome comparisons. Genomes assembled using paired-end DNA-seq data from male and female *T. ni* pupae are compared with the Hi5 genome as the reference. The dot plots show genome alignments for contigs ≥1 kb. (E) Numbers of genes in lepidopteran genomes. (F) Positions of telomeric repeats: position of (TTAGG)_n_ longer than 100 nt. (G) Transposons in *T. ni* subtelomeric regions. (H) Repeat statistics for the *T. ni* genome. (I) Transposon family divergence rates.

**Supplementary file 2**. Genes encoding small RNA pathway proteins. (A) Genes encoding miRNA and siRNA pathway proteins. (B) Genes in encoding piRNA pathway proteins.

**Supplementary file 3**. *T. ni* miRNAs, siRNAs and piRNAs. (A) miRNA annotation. (B) Mapping statistics for endogenous siRNAs in *T. ni* and *D. melanogaster*. (C) piRNA cluster lengths. piRNA cluster coordinates in Hi5 (D), ovary (E), testis (F), female thorax (G), and male thorax (H).

**Supplementary file 4**. mirDeep2 output for *T. ni* miRNAs

**Supplementary file 5**. Genomes used in this study.

**Supplementary file 6**. *T. ni* detoxification-related genes. (A) P450 gene counts by clade in *T. ni* and *B. mori*. (B) Sequences of P450 proteins. (C) Sequences of glutathione-*S*-transferase proteins. (D) Carboxylesterase gene counts by clade in *T. ni* and *B. mori*. (E) Sequences of carboxylesterase proteins. (F) ATP-binding cassette transporter gene counts by clade in *T. ni* and *B. mori*. (G) Sequences of ATP-binding cassette transporter proteins.

**Supplementary file 7**. *T. ni* chemoreception genes. (A) Sequences of olfactory receptor proteins. (B) Sequences of gustatory receptor proteins. (C) Sequences of ionotropic receptor proteins.

**Supplementary file 8**. Genes in the juvenile hormone biosynthesis and degradation pathways.

**Supplementary file 9**. Primers and sequences.

**Supplementary file 10**. Single-stranded DNA donor purification

